# Condensin II mediates efficient chromatid resolution and resistance to genotoxic stress in Arabidopsis

**DOI:** 10.1101/2025.01.21.634090

**Authors:** Jovanka Vladejić, Klara Prochazkova, Eva Dvořák Tomaštíková, Zynke de Cock, Jana Zwyrtková, Kateřina Kaduchová, Markéta Pavlíková, Ales Pecinka

## Abstract

Genome functions are regulated by the means of chromatin and 3D chromosome organization. Dynamic condensation and relaxation of chromosomes during cell cycle is largely controlled by the Condensin complexes. We mapped mutants in Condensin II subunits *SMC2A*, *CAP-D3* and *CAP-H2* as hypersensitive to DNA-protein crosslink (DPC) inducer zebularine. This suggested that Condensin II complex is required for resistance to genotoxic stress in Arabidopsis and prompted us to explore the underlying phenotypes. We show that the Condensin II role in resistance to zebularine is independent of the DNA damage response signaling by SOG1 and the homology-directed repair pathway. Furthermore, we found that Arabidopsis Condensin II mutants have incompletely condensed mitotic chromosomes and show abnormal chromatin connections during anaphase. Based on the structural and temporal analyses, we propose that the connections represent catenated chromatids. The chromosome connections were more frequent when the Condensin II mutants were treated by zebularine or TOP2 crosslinker and inhibitor ICRF-187. Altogether, we demonstrate that the proper large-scale chromatin organization by Condensin II is important for resistance to DNA damage response-inducing agents in Arabidopsis.

## INTRODUCTION

Maintenance of chromosome integrity and a successful passage of chromosomes through cellular generations require dynamic organization at multiple levels (Fraser et al., 2015; Dixon et al., 2016). Key components performing full-scale chromosome organization are STRUCTURAL MAINTENANCE OF CHROMOSOMES (SMC) complexes (Uhlmann, 2016). The eukaryotic SMC complex core is formed by a heterodimeric pair of SMC proteins and additional complex-specific subunits with catalytic and/or structural functions (Haering and Gruber, 2016). The Cohesin complex is responsible for associations of sister chromatids from the S-phase until the end of anaphase across phylla, and data from animals suggest that also for the formation of transcription regulatory loops during interphase (reviewed in e.g. Nasmyth and Haering, 2009). The SMC5/6 complex is widely involved in DNA damage repair (reviewed in e.g. Díaz and Pecinka, 2018; Roy et al., 2024), where it is necessary for normal levels of homology-directed repair (HDR), in Arabidopsis possibly via an interaction with the Cohesin complex (Watanabe et al., 2009), and SUMO-dependent repair (Dvořák Tomaštíková et al., 2023). Lastly, the Condensin complex mediates chromosome condensation during nuclei divisions and biophysical studies using yeast Condensin suggested asymmetric DNA loop extrusion activity through the SMC ring (Ganji et al., 2018).

Plants, similarly to metazoa, contain two Condensin complexes I and II. They share the same core subunits SMC2 and SMC4, but differ by the CONDENSIN-ASSOCIATED PROTEIN (CAP) subunits (Hirano, 2012). In *Arabidopsis thaliana* (Arabidopsis), the core subunits are represented by two paralogs of SMC2 (SMC2A and SMC2B) and three paralogs of SMC4 (Schubert, 2009). Condensin I-specific subunits include CAP-D, CAP-G and CAP-H while Condensin II is associated with CAP-D3, CAP-G2 and CAP-H2 subunits (Schubert, 2009). The principal difference between the two Condensin complexes is their localization during interphase and temporal dynamics of their loading onto chromosomes (Fujimoto et al., 2005). During interphase, Condensin II is present in the nucleus and thus starts the chromosome condensation and Condensin I that is located in cytoplasm reaches chromosomes only after the dissolution of the nuclear membrane inprophase. In Arabidopsis, Condensin II complex is required for normal chromosome function during mitosis and meiosis (Smith et al., 2014; Sakamoto et al., 2019, 2022), stress resistance (Sakamoto et al., 2011) and seed development (Liu et al., 2002; Siddiqui et al., 2003, 2006). Arabidopsis nuclei have specific 3D interphase chromosome distribution of heterochromatic chromocenters (CCs) and corresponding chromosome territories (Fransz et al., 2003; Pecinka et al., 2004). This pattern is greatly modified in Condensin II mutants where the CCs cluster together (Sakamoto et al., 2019; Municio et al., 2021). The establishment of proper 3D interphase genome organization is a result of cooperation between Condensin II and the nuclear membrane-embedded LINC complex (Sakamoto et al., 2022). Furthermore, Condensin II is needed for plant resistance against boron toxicity because CAP-G2 and CAP-H2 subunits were mapped in a genetic screen for sensitivity to excessive boron (Sakamoto et al., 2011). Based on the upregulated DNA damage repair (DDR) genes in *cap-g2* and *cap-h2* plants and their sensitivity to DNA double-strand break (DSB)-inducing agents, it was proposed that Condensin II plays a role in the maintenance of plant genome stability. However, the underlying mechanism remains unclear.

Genome stability is maintained by an intricate network of DDR pathways consisting of modules specialized for DNA damage recognition, signaling and, ultimately, repair (Hu et al., 2016). To study the mechanisms of maintenance of plant genome stability, we developed a system using cytotoxic non-methylable cytidine analog zebularine. Zebularine was primarily described as an epigenetic inhibitor leading to reduced DNA methylation and CC decondensation (Baubec et al., 2009). However, our recent data show that it also affects genome stability in Arabidopsis by causing enzymatic DNA protein crosslinks (DPCs) with DNA METHYLTRANSFERASE 1 (MET1) (Prochazkova et al., 2021). DPCs represent a type of DNA damage where proteins are covalently bound to DNA and form a physical barrier for DNA processing enzymes such as DNA and RNA polymerases (Stingele et al., 2017). At cytological resolution, zebularine-induced DPCs accumulate at the heterochromatic fraction of *45S rDNA* CCs. To uncover the molecular pathways involved primarily in the repair of DPCs, we set up a forward-directed genetic screen named HYPERSENSITIVE TO ZEBULARINE (HZE). The mapping of *HZE1* as *SMC6B* and subsequent analysis pointed to the important role of the SMC5/6 complex and its SUMOylation activity in the repair of zebularine-induced DPCs at heterochromatic loci (Dvořák Tomaštíková et al., 2023). Furthermore, the finding of RAD3-like helicase *REGULATOR OF TELOMERE ELONGATION 1* (*RTEL1/HZE2*) and Polymerase theta *TEBICHI* (*TEB/HZE3*) indicated interplay of HDR in detoxification from heterochromatic DPCs in Arabidopsis (Dvořák Tomaštíková et al., 2024).

Here, we present the mapping and characterization of three new *HZE* complementation groups corresponding to various subunits of the Condensin II complex and representing a functionally distinct class of *HZE* mutants. We show that the sensitivity of Condensin II to zebularine and TOPOISOMERASE 2 inhibitor ICRF-187 is associated with defects in chromosome condensation and sister chromatid resolution during mitotic anaphase.

## RESULTS

### *HZE4* encodes Condensin complex subunit *SMC2A*

The 20 µM zebularine-treated *hze4-1* plants showed 36.9 ± 2.6 % root length compared to mock-treated plants (Figure 1a,b). This was a significantly greater reduction compared to the 49.9% for wild-type (WT) but significantly less than for the sensitive control *smc6b-1* plants with only 10.2 ± 0.1 % of root length (ANOVA with Tukey, *P < 0.05*).

**Figure 1.**
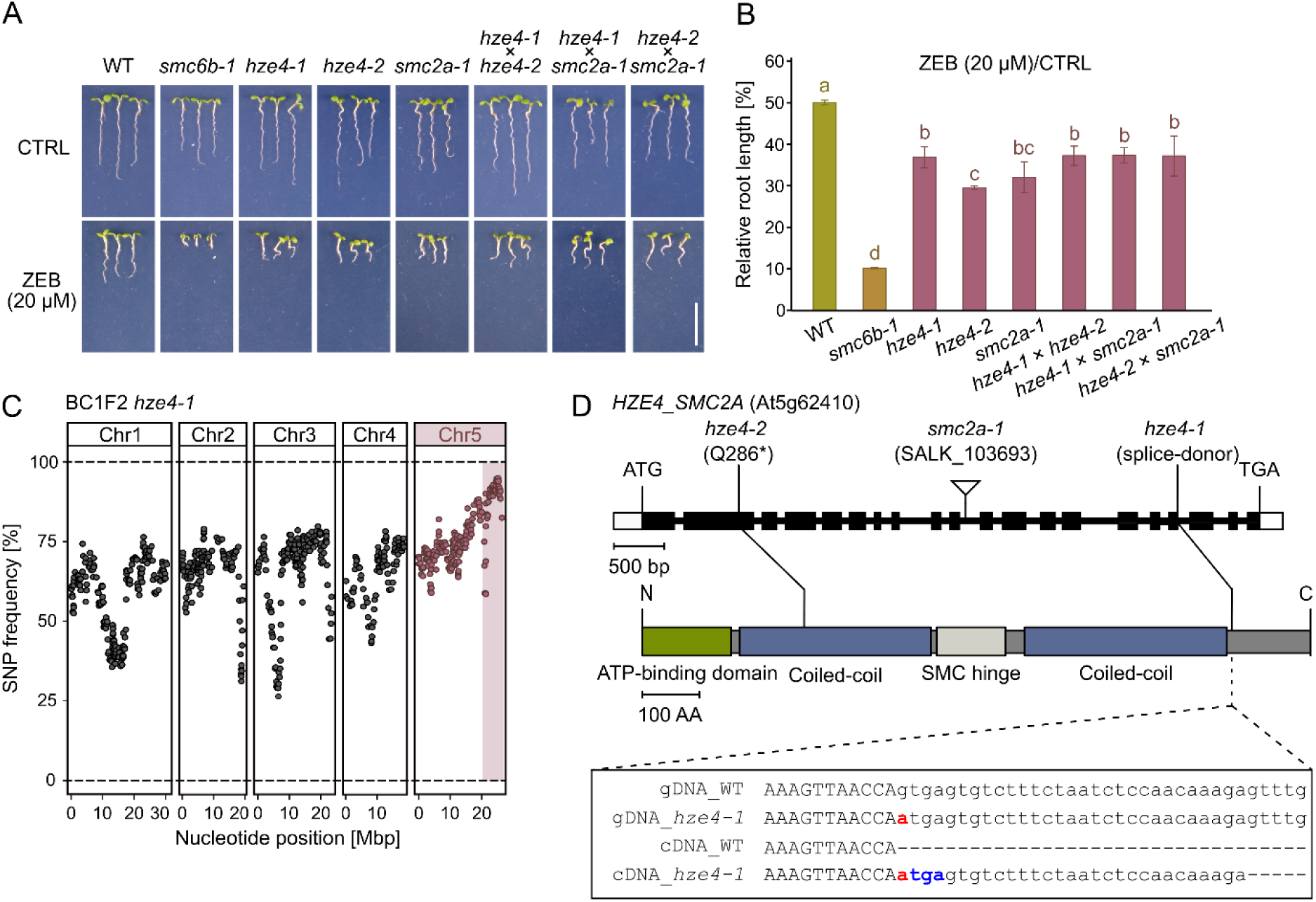
HZE4 encodes Condensin core subunit SMC2A. (a) Representative seven-day-old plants grown on control (CTRL) and 20 μM zebularine-containing (ZEB) ½ MS media. Scale bar, 1 cm. (b) Relative root length of seven days-old zebularine-treated plants. Error bars are the standard deviation of the means of three biological replicates (14-25 plants per replicate). Genotypes marked with the same letter did not differ (P < 0.05) according to one-way ANOVA followed by Tukey’s test. (c) Identification of the chromosomal region containing hze4-1 mutation in zebularine-sensitive plants selected from F2 hze4-1 × WT segregating population. The dots represent the mean frequency of five consecutive high-confidence SNPs. The candidate region on chromosome 5 is highlighted by a light red background. (d) The models of the HZE4 gene and protein with indicated mutations. The inset shows HZE4 wild-type and hze4-1 genomic DNA (gDNA) and copy DNA (cDNA) sequences. Exonic and intronic sequences are in the upper and lower case letters, respectively. The causal mutation is in red and the premature stop codon is in blue.

We identified the candidate causal mutation in *hze4-1* using mapping-by-sequencing (MBS). The main candidate region was at the end of the bottom arm of chromosome 5 (Figure 1c, Figure S1a). However, only the SNP position chr5:25,061,626 was predicted as a high-effect mutation, affecting a splice-donor site at the beginning of intron 17 in gene At5g62410, corresponding to *SMC2A* (Figure 1d; Figure S1b). We sequenced the cDNA and found that the exon 17 was extended for 29 bp, causing a frameshift leading to a premature STOP codon and presumably a truncated SMC2A protein lacking the C-terminus (Figure 1d, inset). During the subsequent screening, we identified *hze4-2* allele, showing a significant 70.5 % root reduction upon 20 µM zebularine treatment (Figure 1a,b). The *hze4-2* plants carried a G→A mutation in exon 2 of *SMC2A*, causing a premature STOP codon (Q286*; Figure 1d, Figure S1c,d). A similar 67.9 % reduction in root length on 20 µM zebularine was found for the T-DNA insertional mutant *smc2a-1* (Figure 1a,b,d). Genetic complementation tests using F1 hybrids between *hze4-1*, *hze4-2,* and *smc2a-1* revealed a similar level of sensitivity (Figure 1a,b), suggesting that all these mutants are allelic.

Hence, *HZE4* corresponds to *SMC2A*, which has been identified as a core subunit of both Condensin I and II complexesin Arabidopsis (Schubert, 2009).

### *HZE5* and *HZE6* link hypersensitivity to zebularine with Condensin II complex

Later, we mapped two more Condensin subunits, named *HZE5* and *HZE6*. The four mutant alleles of *HZE5* (*hze5-1* to *hze5-4*) showed 62.9 %, 73.4 %, 75.3 %, and 76.8 %, respectively, reduction of root length in response to 20 μM zebularine (Figure 2a,b). The candidate region of all *hze5* lines mapped to the bottom arm of chromosome 4 and gene At4g15890 corresponding to *CAP-D3* (Figure 2c; Figure S2, S3, S4, and S5). An interesting technical aspect of the mapping was that the relative root lengths of *hze5-1* and WT plants were not significantly different at 95% probability (*P < 0.05*), yet, we were able to map this mutant using MBS (Figure 2c; Figure S2). This demonstrates the power of MBS-based forward genetics, where even small differences can lead to accurate identification of the causal loci (Dvořák Tomaštíková and Pecinka, 2025). The *hze5-1* had a G→A transition in exon 2 causing G631D mutation (Figure S2c). The *hze5-2* and *hze5-3* contained C/G→T/A transitions leading to Q1045* and W656* mutations, respectively. The *hze5-4* line carried G→A transition in the splice-donor site of intron 9 leading to two alternative splice variants (Figure 2c, Figure S3c, S4c and S5c). The *hze5-4a* transcript retained intron 9 and had a premature STOP codon following amino acid 1104, while the *hze5-4b* transcript lost five amino acids causing a premature STOP codon after amino acid 1131 (Figure 2c). The sensitivity assays using the *cap-d3-2* T-DNA mutant and complementation tests using F1 crosses of *hze5-1* to *5*-*4* alleles with *cap-d3-2* revealed hypersensitivity to zebularine, confirming that the *hze5* mutations are allelic to *cap-d3* (Figure 2a).

**Figure 2.**
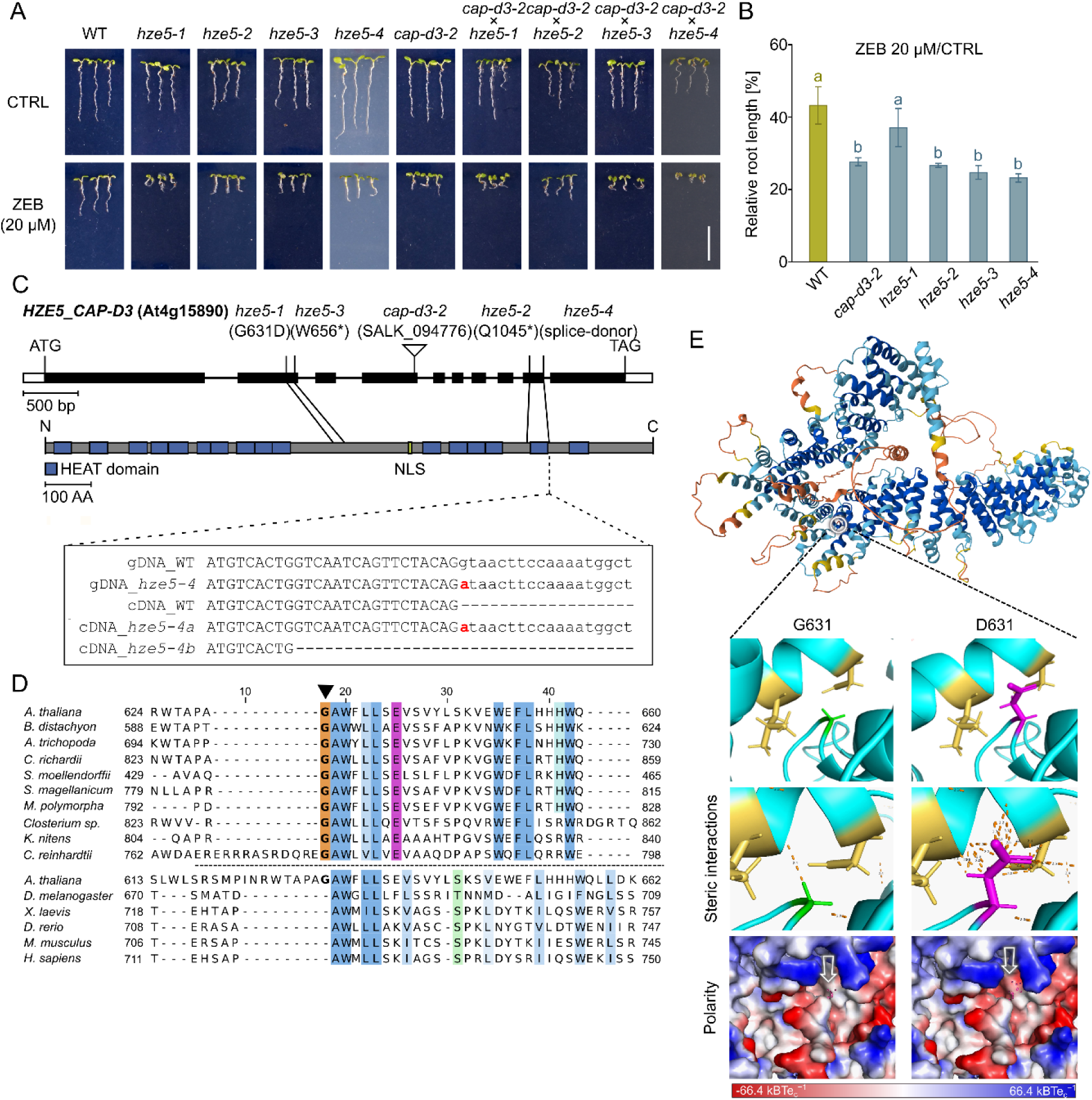
HZE5 encodes Condensin II subunit CAP-D3. (a) Representative seven-day-old plants grown on media without (CTRL) or with 20 μM zebularine (ZEB). Scale bar, 1 cm. (b) Relative root length of seven days-old zebularine-treated plants. Error bars are the standard deviation of the means of three biological replicates (14-25 plants per replicate). Genotypes marked with the same letter did not differ (P < 0.05) in Tukey’s test. (c) The models of the HZE5 gene and protein showing locations of the mutations. The inset shows the impact of the splice variant mutant hze5-4 on the mature transcript. Exonic and intronic sequences are in the upper and lower case letters, respectively, and the causal mutation is marked in red. (d) Multiple protein alignment of the CAP-D3 region containing G631D mutation (in bold and marked by the arrowhead). The Arabidopsis sequence was aligned with sequences of other plant species in the upper part, and with other model organisms in the bottom part. (e) Alpha-fold-based model of Arabidopsis CAP-D3 with a detailed view on the wild-type and hze5-1 mutated amino acid 631.

The most interesting HZE5 mutation was G631D substitution in *hze5-1*. It did not affect any of the 16 annotated HEAT domains (Figure 2c), but the plants had strongly reduced root length (Figure 2a), suggesting that it affected a functionally important residue. We performed alignment of CAP-D3 from multiple species and found that G631 was conserved among all tested plants but relatively variable in animals (Figure 2d). This amino acid is adjacent to a AWXXLXE^631-637^ motif conserved in all analyzed plant species. Inspection of the predicted protein structure suggested that Arabidopsis G631 is the first amino acid before an alpha-helix that is embedded deep in the protein and surrounded by several heat domains (Figure 2e). Modeling of the mutated protein (G631D) showed an more negative charge at the mutated region and a possible sterical hindrance that could affect protein folding (Figure 2e, bottom panel).

The *hze6-1* and *hze6-2* mutants showed 63.1 % and 71.3 % root length reduction after 20 μM zebularine treatment, respectively (Figure 3a,b). The causal mutations were mapped to the top arm of chromosome 3 where both lines carried high-effect mutations in At3g16730 corresponding to *CAP-H2* (Figure S6 and S7). Both alleles contained G→A transitions causing loss of splice-donor site in *hze6-1* and a premature stop W359* in *hze6-2* (Figure 3c). We confirmed sensitivity to 20 μM zebularine using *cap-h2-2* T-DNA mutant plants and F1 hybrids (Figure 3a,b), confirming that *HZE6* is allelic with *CAP-H2*.

**Figure 3.**
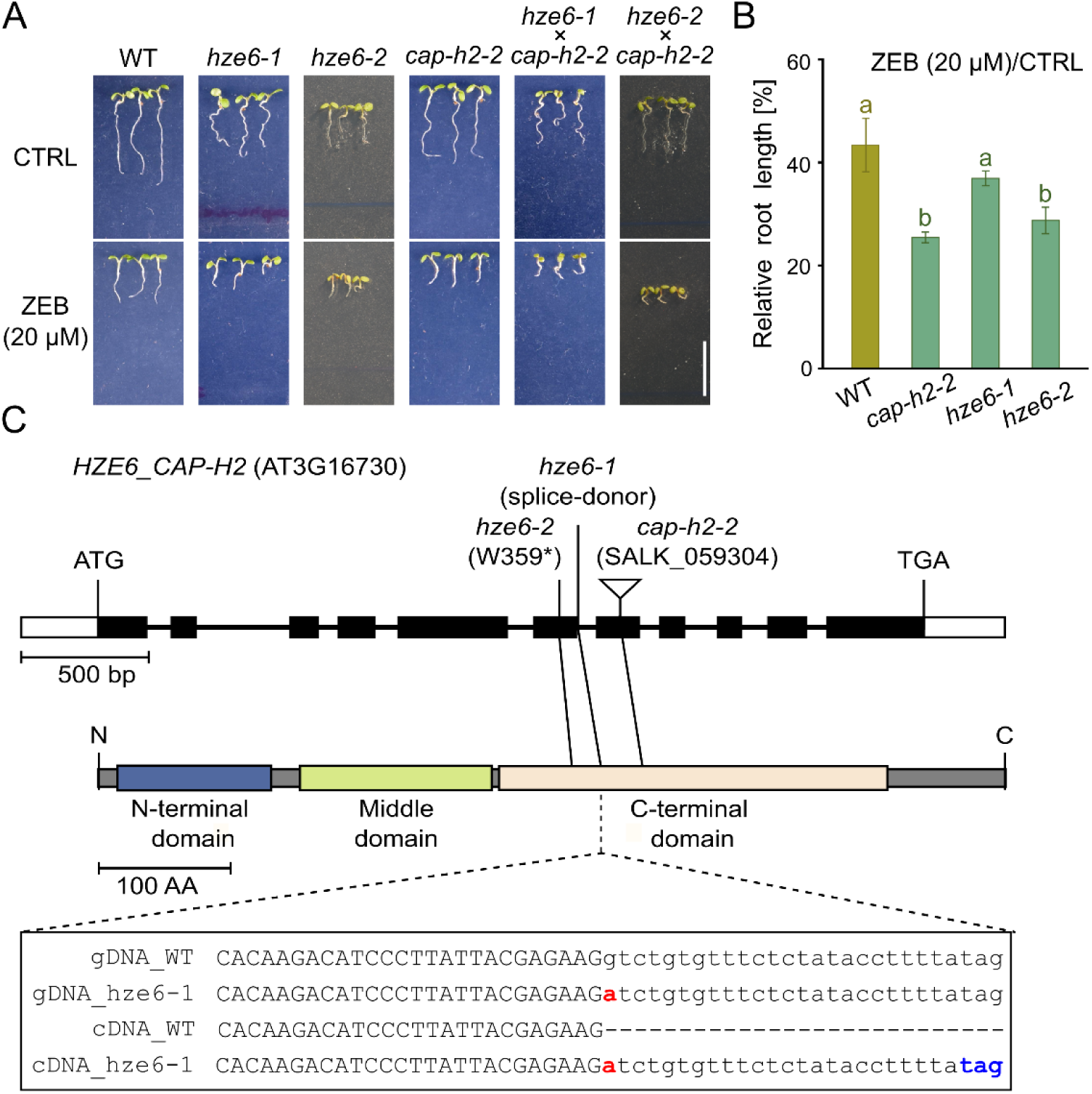
HZE6 encodes Condensin II subunit CAP-H2. (a) Representative seven-day-old plants grown on media without (CTRL) or with 20 μM zebularine (ZEB). Scale bar, 1 cm. The last two columns represent F1 hybrid phenotypes of the complementation test crosses. (b) Relative root length of seven days-old zebularine-treated plants. Error bars are the standard deviation of the means of three biological replicates (14-25 plants per replicate). Genotypes marked with the same letter did not differ (P < 0.05) in Tukey’s test. (c) The models of the HZE6 gene and protein showing locations of the mutations. The inset shows the impact of the splice variant mutant hze6-1 on the mature transcript. Exonic and intronic sequences are in the upper and lower case letters, respectively, the causal mutation is marked in red and the premature stop codon in blue.

Hence, we conclude that HZE5 encodes CAP-D3, the HEAT-A subunit of Condensin II, and HZE6 is CAP-H2, the β-kleisin subunit of the same complex, suggesting that Condensin II is required for resistance to zebularine toxicity.

### Condensin II mutants are sensitive to TOPOISOMERASE 2 inhibitor ICRF-187

To explore the extent of the role of Condensin II in DDR, we exposed *cap-d3*, *cap-g2* and *cap-h2* plants to chemicals inducing various types of DNA damage (Figure 4; Figure S8). We confirmed previously observed sensitivity to radiomimetic agents bleocin and zeocin (Sakamoto et al., 2011). Condensin II mutants showed hypersensitivity to DNA inter-strand crosslinker mitomycin C but not to camptothecin, an enzymatic crosslinker of TOPOISOMERASE 1 (TOP1). Surprisingly, treatment with ICRF-187 caused 87-92 % root length reduction for Condensin II mutants compared to 7 % reduction for WT plants (Figure 4a,b). ICRF-187 stabilizes TOP2 on a DNA molecule at the end of its enzymatic cycle, causing its simultaneous inhibition and formation of an enzymatic DPC (Nitiss, 2009).

**Figure 4.**
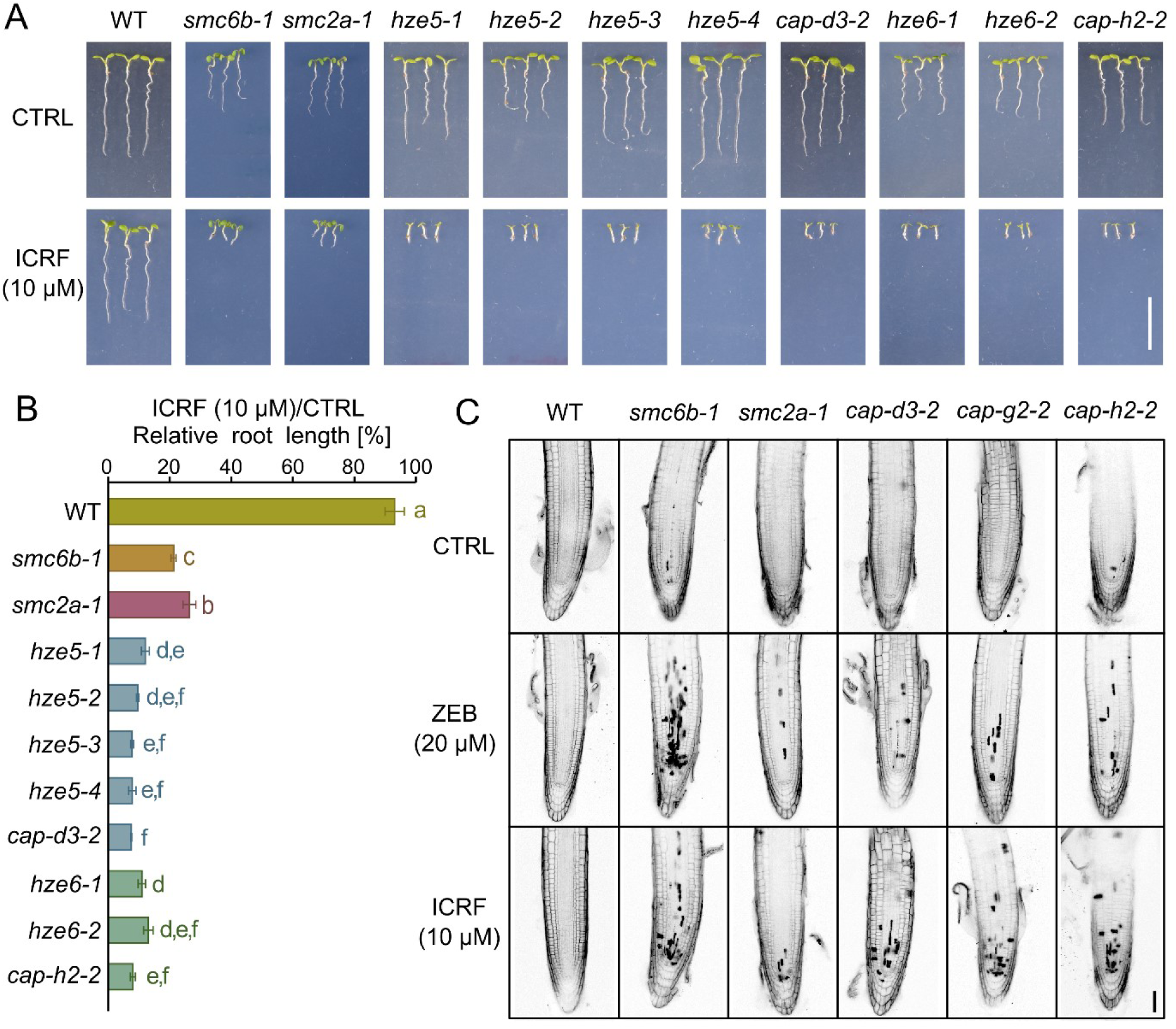
Condensin II mutants are hypersensitive to TOP2 inhibitor ICRF-187. (a) Representative seven-day-old plants grown on media without (CTRL) or with 10 μM ICRF-187 (ICRF). Scale bar, 1 cm. (b) Relative root length of seven days-old zebularine-treated plants. Error bars are the standard deviation of the means of three biological replicates (14-25 plants per replicate). Genotypes marked with the same letter did not differ (P < 0.05) in Tukey’s test. (c) Cell death analysis using propidium iodide staining after 24-hour CTRL, ZEB or ICRF treatments. Scale bar, 50 µm.

Using propidium iodide staining assays, we tested whether the reduced root growth in Condensin II mutants coincides with an increased frequency of cell death (Figure 4c). Indeed, there was a moderate amount of dead cells in Condensin II mutants after zebularine and ICRF-187 treatments compared to a low number in WT and high number in the control *smc6b-1* plants.

Thus, Condensin II is required for plant resistance to structurally different types of DNA damage. The greatest sensitivity was observed for zebularine and the TOP2 inhibitor ICRF-187.

### Condensin II acts independently of canonical DDR

A large part of Arabidopsis DDR is orchestrated by the ANAC-type transcription factor SUPPRESSOR OF GAMMA RADIATION 1 (SOG1) (Bourbousse et al., 2018; Ogita et al., 2018). Therefore, we tested for a dependence of Condensin II function on the SOG1 signaling by analysis of double mutants. The mean root length of mock-treated *cap-g2-2* (12.8 mm) and *cap-h2-2* (11.2 mm) plants were comparable to WT plants (13.9 mm) and the roots of *sog1-1* were longer (17.5 mm) (Figure 5a,b; Figure S9a). The roots of *cap-g2-2 sog1-1* and *cap-h2-2 sog1-1* plants reached only 5.5 and 6.7 mm, respectively, suggesting a synergistic interaction and indicating that Condensin II and SOG1 function independently under ambient conditions. The same conclusion held true also after testing the double mutants on 20 μM zebularine or 1 μM ICRF-187. We also observed increased cell death in the double mutant plants, which concentrated atypically in the root elongation zone (Figure 5c; Figure S9a).

**Figure 5.**
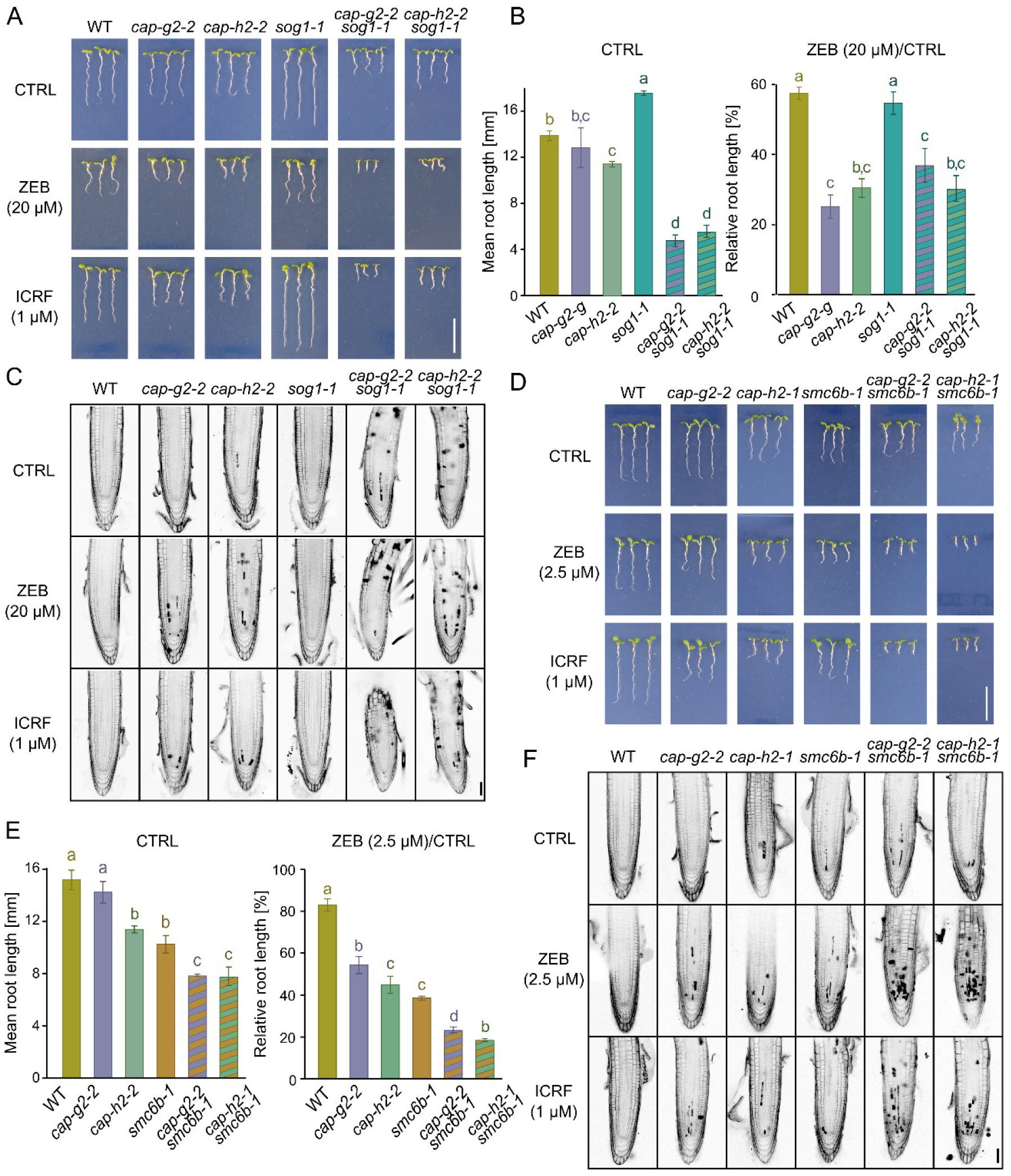
Condensin II functions independently of SOG1 and SMC5/6 complex pathways. (a) Analysis of genetic interaction between Condensin II and SOG1. Representative seven-day-old plants grown on control media (CTRL) or media supplemented with 20 μM zebularine (ZEB) or 1 μM ICRF-187 (ICRF). Scale bar, 1 cm. (b) Absolute and relative root length of seven days old zebularine-treated plants. Error bars are the standard deviation of the means of three biological replicates (14-25 plants per replicate). Genotypes marked with the same letter did not differ (P < 0.05) in Tukey’s test. The ICRF data are provided in Figure S9A. (c) Cell death analysis using propidium iodide staining after 24 hours of CTRL, ZEB or ICRF treatments. Scale bar, 50 µm. (d) and (e) Analysis of genetic interaction between Condensin II and SMC5/6 complex. The experiment was performed as described in (a) and (b) but with a lower concentration of zebularine (2.5 μM) to be able to assess possible additive/synergistic effects in the double mutant plants. Quantitative data for ICRF are provided in Figure S9B. (f) Cell death analysis performed as described in (c).

Furthermore, we tested for the crosstalk between Condensin II and SMC5/6 complex that had a prominent role in repair of zebularine-induced DPCs (Dvořák Tomaštíková et al., 2023). We developed *cap-g2-2 smc6b-1* and *cap-h2-1 smc6b-1* double mutant plants. These showed significantly reduced root length compared to the single mutant and WT plants under mock and 2.5 µM zebularine and 1 µM ICRF-187 treatments (Figure 5d,e; Figure S9b). The cell death assays in genotoxin-treated double mutant plants supported the outcome of the root length assays by revealing an additive phenotype compared to the parental lines with dead cells accumulating in the meristematic zone (Figure 5f).

These results suggest that Condensin II acts independently of SOG1 and SMC5/6 complex.

### Condensin II mutants have WT-like frequency of HDR and genome mutation rate

To directly test for the role of Condensin II in HDR, we generated *cap-g2-2* and *cap-h2-1* carrying single-strand annealing (SSA) HDR traps 651 and B11 (Puchta et al., 1995). The mutant plants showed WT-like SSA frequencies under control conditions (Figure S10). Subsequently, we applied low doses of zebularine (2.5 μM) and ICRF-187 (2.5 μM) to stimulate DDR but to avoid extensive cell death. Zebularine-treated B11 line showed a 1.7-fold increased SSA while this trend was significantly higher (3.7 and 6.9-fold) in *cap-g2-2* and *cap-h2-2* plants, respectively (Figure S10a). This suggested that Condensin II might be a SSA repressor. However, the following experiments with zebularine-treated 651 line and ICRF-treated B11 and 651 lines (Figure S10) showed either no differences or even significant SSA reduction in the mutants, often due to the presence of one or two plants with marginal values. These overall minor differences likely represent inherent variation typical for these HDR reporter lines rather than a uniform mutant effect.

The SSA trap lines represent an artificial recombination substrate. Therefore, we aimed to estimate the mutation rate of Condensin II mutants at a genome-wide scale and in natural sequences similarly to previous studies (Ossowski et al., 2010; Willing et al., 2016). To this end, we propagated WT and *cap-d3-2* plants for five generations, three siblings per genotype were whole genome sequenced, and novel mutations were identified. we found a similar numbers of three and five single nucleotide polymorphisms (SNPs) in WT and *cap-d3-2* plants, respectively, and no insertions and/or deletions were found in any of the plants (Table S1).

We conclude that Condensin II complex is not involved directly in HDR by SSA and that its mutants do not suffer from increased mutation rate.

### Condensin II mutants show abnormalities at mitotic anaphase

Next, we hypothesized that the mutant sensitivity to genotoxic stress might be connected with mitotic chromosome structure and/or dynamics. We first examined mitotic anaphase figures (Figure 6a). WT plants contained 92.7 % regularly looking anaphases with two groups of well-condensed and clearly separated sister chromatids without any microscopically observable chromatin connections and/or laggards (Figure 6b). On the contrary, there were only 38.1 % regularly-looking anaphases in *cap-g2-2* and 29.2 % in *cap-d3-*2 plants. The abnormalities included (i) incompletely condensed chromosome arms and (ii) chromatin fibres connecting separating chromosome complements (Figure 6a,b). After the treatment with 40 μM zebularine, the frequency of normally-looking anaphases decreased from to 84 % in WT, 24.3 % in *cap-d3-2* and 25.7 % in *cap-g2-2*. The effect of 10 μM ICRF-187 treatments was even stronger, and we found 64 % normal anaphases in WT, 0 % in *cap-d3-2* and 3 % in *cap-g2-2*. Interestingly, the mitotic defects observed in Condensin II mutants and/or drug-treated plants included incompletely condensed chromosomes visible as parts of chromosome arms protruding from the bulk of chromomes during anaphase. Futhermore, there were chromatin fibers connecting separating sets of chromatids. Although these structures resembled anaphase bridges, we did not observe any lagging chromosome fragments - a typical byproduct of reciprocal translocations and formation of dicentrics. This suggests that the chromatin connections might represented an earlier mitotic stage represented by the incompletely condensed chromosome arms.

**Figure 6.**
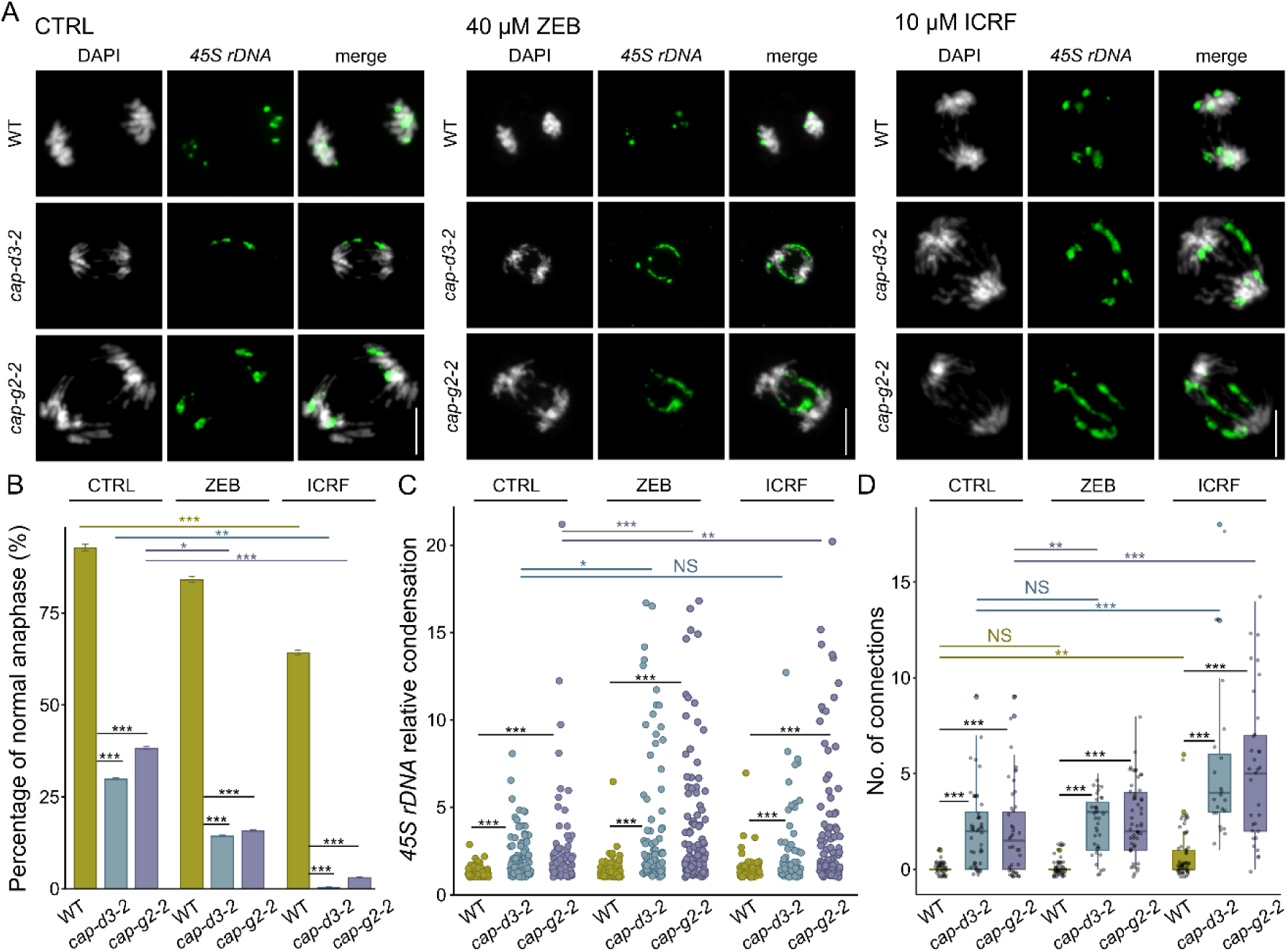
Condensin II mutants show chromatid entanglements. (a) Representative mitotic anaphases from inflorescences of wild-type (WT) and Condensin II mutants without (CTRL) and after 40 μM zebularine (ZEB) and 10 μM ICRF-187 (ICRF) treatments. The 45S rDNA (green) was labeled by fluorescence in situ hybridization (FISH) and DNA was stained with DAPI (grey). Bar, 5 μm. (b) Normally looking anaphases in WT and Condensin II mutants. In total 23 to 63 anaphases per experimental point were scored. Error bars represent a confidence interval for each value. The statistical relevance of the results was calculated in the Chi-square test. Only the significant differences are indicated in the graph, all values can be found in Table S3., * P < 0.05, ** P < 0.01, *** P < 0.001. (c) Relative condensation of 45S rDNA signals (FISH signal length/width) at anaphase. Each dot on the jittered strip plot represents one value. Statistical significance was tested with the Kruskall-Wallis H-test with post hoc Conover-Iman test of multiple comparisons using rank sums with Benjamini-Hochberg procedure (P < ½ α, α = 0.05) in R 4.2.1. For a full list of values see Table S4. NS – not significant. (d) Number of chromatin connections between separating chromosome complements (including those without 45S rDNA). Box plots show the individual numbers of connections in each sample as grey dots. The hinges are in the 1st and 3rd quartile, with a marked median. Whisker marks show the lowest or highest value within the 1.5 interquartile range below or above hinges. For the full list of statistical significance see Table S5.

To support this empirical observation with quantitative data, we measured the relative compaction of *45s rDNA* loci, which represent a well-defined, highly compact heterochromatic repeat arrays on the top arms of chromosomes 2 and 4 (Arabidopsis Genome Initiative, 2000). We defined the condensation as the average *45S rDNA* fluorescence *in situ* hybridization (FISH) signal length/width (L/W; Figure 6c). The *45S rDNA* loci had a low average L/W ratio 1.28 at anaphase of WT plants. The 40 μM zebularine and 10 μM ICRF-187 treatments had only a minor effect on the average L/W in WT (1.37 and 1.46, respectively) but in a few cases, we observed also strongly decondensed *45S rDNA* arrays (maximal L/Ws = 6.47 and 6.96, respectively). The *cap-d3-2* and *cap-g2-2* mutations led to approximately doubled average L/W ratio (2.12 and 2.36, respectively), indicating incomplete *45S rDNA* condensation. We also found several cases with *45S rDNAs* appearing as fine threads connecting separating sister chromatids (maximum L/Ws 8.07 and 21.19, respectively). The combination of mutations and drug treatments increased the average L/W there was a high number of events with an extreme decondensation (L/W >10) (Figure 6c).

We also quantified the number of chromatin connections per anaphase which reached 0.07 for WT, 2.08 for *cap-d3-2* (29.7-fold increase) and 2.07 (29.6-fold increase) for *cap-g2-2* (Figure 6d). The zebularine and ICRF-187 treatments increased the frequency of connections per WT anaphase to 0.16 and 0.774 (2.3- and 11.1-fold increase, respectively). In *cap-d3-2* and *cap-g2-2* plants, there were 2.31 and 2.69 connections per anaphase after 40 μM zebularine (1.1- and 1.3-fold increase over mock-treated mutants, respectively), and 5.30 and 5.30 connections after 10 μM ICRF-187 treatments (2.5- and 2.6-fold increase over mock-treated mutants, respectively). Interestingly, although the connecting chromatin fibers regularly involved *45S rDNAs*, they did not contain centromeric repeats (not shown). This suggested that the problems affected mainly the telomere-proximal parts of the chromosomes. Importantly, we did not observe any lagging chromosomes or chromosome fragments in mitoses of the *cap* mutant plants excluding reciprocal translocation and formation of dicentric chromosomes as the causal effect. Although this was a surprising finding, it indicates that the sensitivity of Condensin II mutants to DNA-damaging agents is not due to an increased genome fragmentation.

Therefore, Condensin II mutants show less condensed mitotic chromosomes and abnormalities during sister chromatid resolution at anaphase that likely represent catenated sister chromatids. The defects are aggravated by the genotoxic stress.

## DISCUSSION

The use of non-methylable cytidine analogs is firmly established as an approach to induce transient DNA demethylation in plant epigenetic research (reviewed in Dvořák Tomaštíková and Pecinka, 2024). We recently demonstrated that the cytidine analog zebularine can be used also for a controlled induction of DPCs in heterochromatic *45S rDNA* regions of Arabidopsis (Prochazkova et al., 2021). The system served as a basis for the forward-directed genetic screen, through which we started to uncover the molecular network involved in the detoxification of zebularine cytotoxic effects in plants. Here, we presented three new complementation groups *HZE4*/*SMC2A*, *HZE5*/*CAP-D3* and *HZE6*/*CAP-H2* that collectively suggest a role of Condensin II complex in ensuring normal resistance to genotoxic stress in Arabidopsis. We show that the involvement of Condensin II is not limited or directly linked to HDR-based DPC repair, but it rather ensures efficient resolution of sister chromatids during anaphase, which seems to be challenged by the complex effect of specific drugs on chromosome structure (zebularine) and/or resolution of catenated DNA (ICRF-187) (Figure 7).

**Figure 7.**
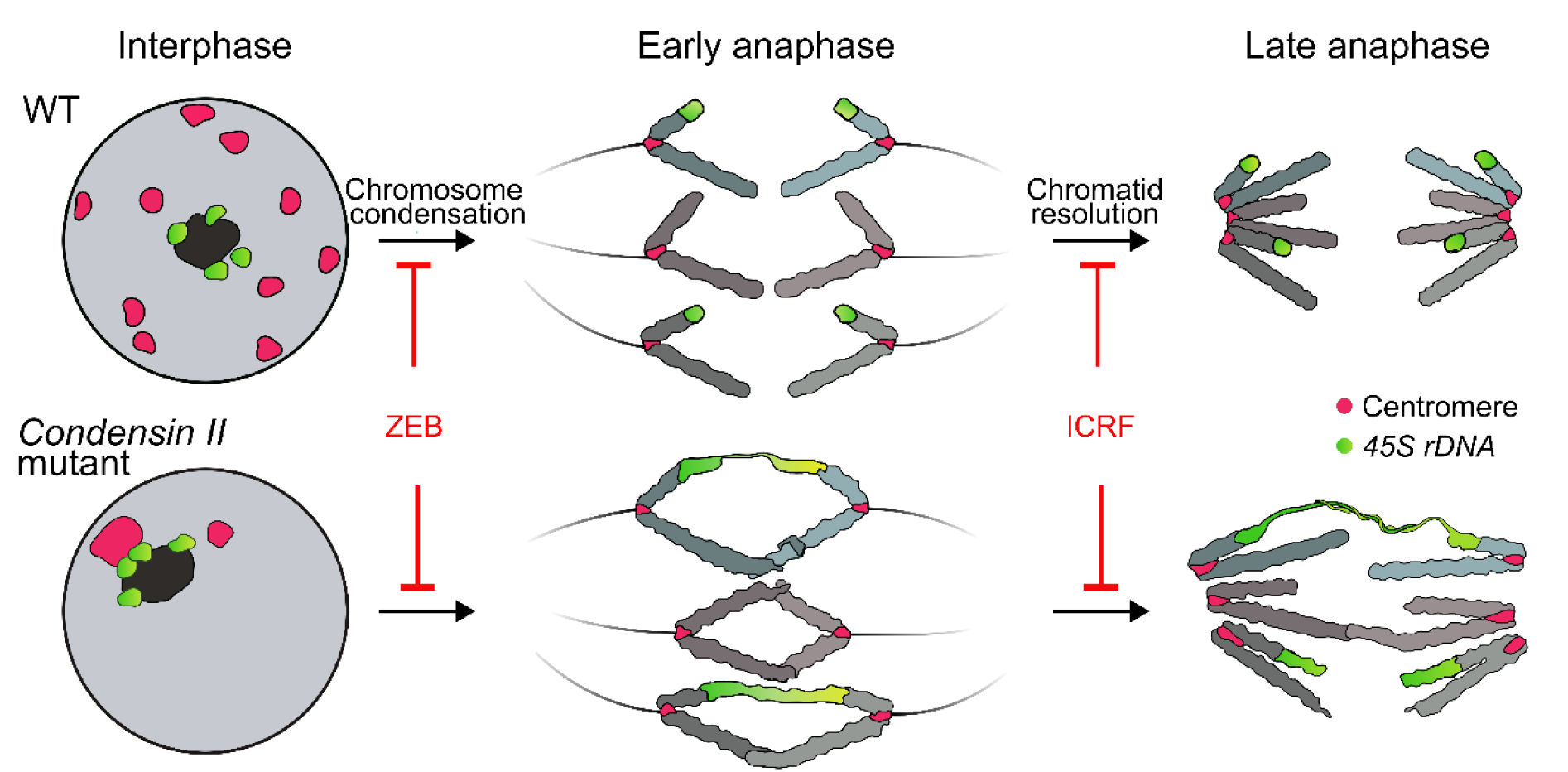
The model of Condensin II effects in sister chromatid resolution during mitotic anaphase in Arabidopsis. Wild-type (WT) interphase nucleus shows regularly distributed centromeric chromocenters (red) and 45S rDNA loci (green) clustered around nucleolus (dark). Normal anaphase is hallmarked by properly condensed chromosomes, compact 45S rDNA loci and absence of chromatin connections. Condensin II mutants show altered interphase chromosome distribution with clustering of centromeric chromocenters (Sakamoto et al., 2019, 2022). During anaphase, separating chromosome complements are often connected by chromatin fibers that regularly include 45S rDNAs and likely represent catenated chromatids. The defects are aggravated by chemical treatments with cytidine analog zebularine and TOPOISOMERASE 2 inhibitor ICRF-187.

Condensin II mutants were previously identified as sensitive to excessive boron which was associated with an increased DDR at the transcriptional level and more fragmented DNA upon zeocin treatment (Sakamoto et al., 2011). However, the underlying mechanism remained unclear. In our study, Condensin II mutants showed moderate to strong sensitivity to wide spectrum of lesions – DPCs (zebularine, ICRF-187 – also TOP2 inhibition), DNA strand breaks (bleocin, zeocin) and DNA inter-strand crosslinks (MMC). The *HZE4*/*SMC2A* mutants had overall lower sensitivity and their phenotypes were more variable, while the sensitivity of mutants in *CAP-D3*/*HZE5*, *CAP-H2*/*HZE6* and *CAP-G2* was stronger and uniform. The lower and heterogeneous sensitivity of *SMC2* is most likely due to its multicopy nature compared to single-copy character of *CAP* genes in Arabidopsis (Siddiqui et al., 2003). Collectively, this points to an important, but not critical, role of Condensin II in plant resistance to genotoxic stress and shows that this complex is unlikely to be limited to a specific type of DNA damage and/or repair pathway. Our experiments indicate that the role of Condensin II in plant DDR might primarily overlap with its function in large-scale chromatin organization and prevention of chromosome entanglements through efficient chromosome condensation.

Genetic analysis of SOG1 and Condensin II double mutants suggested that Condensin II acts independently of the key transcriptional regulator of DNA damage responses SOG1 (Yoshiyama et al., 2009; Bourbousse et al., 2018; Ogita et al., 2018). This is consistent with our finding that HDR by SSA is generally unaffected in Condensin II mutants, except for increased HDR in zebularine-treated line B11. It remains open whether the B11 locus is heterochromatic due to its multi-copy nature (Puchta et al., 1995), and thus reflects a combined effect of zebularine on DNA methylation, heterochromatin compaction, and genome stability (Baubec et al., 2009; Pecinka et al., 2009; Nowicka et al., 2020). Previously, we showed that DDR signaling of zebularine-induced DPCs is mediated redundantly by the ATM and ATR kinases and, in parallel, also by SUMOylation via SMC5/6 complex (Liu et al., 2015; Dvořák Tomaštíková et al., 2023). The analysis of Condensin II and SMC5/6 double mutants revealed that both SMC complexes work independently in the DPC repair. Therefore, we propose that Condensin II-mediated tolerance to genotoxic stress is not coupled with canonical HDR parhways.

Condensin II is localized on chromatin during interphase and starts chromosome condensation during nuclear division (Fujimoto et al., 2005). Furthermore, it facilitates the distribution of (peri)centromeric chromocentres across the nuclear envelope, via interaction with nuclear lamina in Arabidopsis (Sakamoto et al., 2022). Loss-of-function from Condensin II leads to the clustering of centromeric CCs to a single or few large heterochromatic domains observable in a microscope, and an increased frequency of contacts between the centromeric regions that was confirmed via chromatin conformation capture methods (Sakamoto et al., 2019, 2022). Such modified organization will alter the overall distribution of repetitive DNA in the cell nucleus, which might affect the choice of the repair substrate. Treatment of Condensin II mutants with radiomimetic agent zeocin caused a higher number of cytologically detectable γ-H2A.X foci in centromeric regions and an increased amount of DNA fragments in tails in comet assays (Sakamoto et al., 2022). Erroneous repair in repetitive DNA may result in a higher frequency of reciprocal translocations leading to dicentric chromosomes and acentric fragments that are usually lagging behind in anaphase and typically get lost. Therefore, we searched for lagging chromosome fragments in mitosis that would indicate frequent chromosome breakage-fusion events in Condensin II mutants. Interestingly, we observed frequent chromatin connections between the separating chromatids, but there were no lagging fragments. This strongly suggests that the chromatin connections were not anaphase bridges. Instead, they likely represented catenated sister chromatids.

Based on our data and studies on non-plant models, we propose a model on how defective Condensin II function and pharmacological interference with chromatin organization by zebularine or ICRF-187 may lead to the observed phenotypes (Figure 7). Following S-phase, newly sysnthesized chromatids are to some extent catenated, which si sometimes called as sister chromatid intertwinings (SCIs) (Jeppsson et al., 2014). During mitosis, chromosomes gradually compact by the activity of Condensin complex(es), which stimulates the activity of TOP2 that resolves the catenations and thus facilitates a normal sister chromatid segregation (Baxter, 2015; Sen et al., 2016). The lower degree of chromosome condensation in Condensin II mutants is probably not sufficientto fully stimulate a TOP2-based SCI resolution (Sen et al., 2016). As a consequence, at least part of the SCIs persist and represent a significant obstacle towards the normal sister chromatid separation. The catenated sister chromatids then appear as chromatin fibers that are eventually passively resolved at late anaphase or telophase. How this triggers DNA damage response is currently unclear. One possible consequence is the breakage of chromosome arms during phragmoplast formation at the end of mitosis (Schubert and Oud, 1997). Although we did not observe such situations, we cannot fully exclude it. Another variant is that the topological stress of catenated DNA leads to unusual structures that are recognized by specific DNA damage sensors that trigger DDR. The third possibility includes a too-long duration of mitosis that interferes with specific mitotic checkpoint(s) leading to slower divisions and in extreme cases cell death.

The zebularine and ICRF-187 could affect the process in two different ways. Zebularine was shown to reduce chromatin compaction (Baubec et al., 2009; Nowicka et al., 2020). In combination with Condensin II mutations, this might additively reduce mitotic chromosome compaction and thus weaken TOP2 activity in resolving catenations during anaphase. The ICRF-187 stabilizes TOP2 on the DNA at the end of its enzymatic cycle (Classen et al., 2003). This leads to the formation of TOP2 DNA protein crosslinks and at the same time a depletion of TOP2 pool. In WT, the lower amount of TOP2 will reduce the potential for SCI resolution during anaphase. A combination of poorly condensed chromosomes in Condensin II mutants together with the ICRF-187 affected TOP2 pool will subsequently cause massive problems in resolving catenated DNA during anaphase as observed in our experiments.

In conclusion, our study suggests that the proper chromosome compaction ensured by the Condensin II complex is required for normal mitotic chromosome segregation in Arabidopsis. Pharmacological interference with chromatin organization by zebularine in combination with the loss of function from Condensin II leads to abnormal progression of mitosis due to non-resolved SCIs. Furthermore, treatment of Condensin II mutants with ICRF-187 inhibitor suppresses essential TOP2 activity hallmarked by a large number of SCIs and mitotic catastrophe. Hence, our study shows that the large-scale chromosome organization is important for normal genome function during genotoxic stress in plants.

## MATERIALS AND METHODS

### Plant materials and cultivation

Homozygous genotypes of *A. thaliana* Columbia-0 (Col-0) were used throughout this study (unless stated otherwise): Col-0, W35 (Willing et al., 2016), *smc6b-1* (SALK_123114C), *smc2a-1* (SALK_103693), *cap-d3-2* (SALK_094776), *cap-g2-2* (SALK_049790; Sakamoto et al., 2011), *cap-h2-1* (EMS; Sakamoto et al., 2011), *cap-h2-2* (SALK_059304; Sakamoto et al., 2011), 651 and B11 (Puchta et al., 1995), IC9C (Molinier et al., 2004). The T-DNA lines were obtained from the European Arabidopsis Stock Centre (NASC) except for the EMS mutants *heb1-1* (*cap-g2*) and *heb2-1* (*cap-h2*) which were kindly donated by Dr. T. Sakamoto. The genotyping primers are provided in Table S2.

Plant cultivation in soil. The seeds were spread on a water-soaked substrate, and plants at a two-leaf stage were singled into 7 × 7 cm pots. The pots were kept in the air-conditioned phytochamber with a long day condition 16 h light, 150 µmol m^-2^ s^-1^ (fluorescent tubes MASTER TL-D 18W/840, Philips), 21 °C, 8 h dark, 19 °C, 60% humidity.

*In vitro* cultivation. The seeds were surface sterilized (5 min 70% ethanol, followed by 6 min in 8% hypochlorite solution containing 1% Tween 20, washed with copious amounts of sterile water), stratified by keeping wet seeds for 2-4 days in the dark at 4°C, sown on ½ Murashige and Skoog (½ MS) media with 0.6% agar without (mock) or with DNA damaging chemicals in concentrations specified in the text (zebularine, Z4775; mitomycin C, M4287; camptothecin, C9911; ICRF-187/dexrazoxane, D1446, all Sigma-Aldrich; and bleocin, 203410, Calbiochem) and grown in a Percival chamber (16 h light, 120 µmol m^-2^ s^-1^, 21 °C, 8 h dark, 21 °C, 100% humidity). For root length assays, seven days old seedlings were carefully pulled out of the agar, stretched on plates with 1.5% agarose and photographed with a D5600 Nikon camera and 80 mm Nikkor objective. Subsequently, the roots were measured using ImageJ software calibrated with an internal size control. The effect of the drug treatment was assessed by relative root length (%), calculated as a drug/mock condition ratio. Every experiment was performed in three biological replicates, each with at least 20 plants.

### Mutant identification and mapping-by-sequencing

The primary mutant candidates were selected from the M_2_ ethyl methanesulfonate (EMS) mutagenized population as described (Dvořák Tomaštíková et al., 2023). Validated M_3_ candidates were crossed with the W35 WT line and backcross F_2_ mapping population was generated. Approximately 600 plants were grown on a plate with ½ MS media containing 20 μM zebularine. Roughly 100 plants with sensitive and uniform phenotypes were selected, bulked, and their gDNA was isolated using Dneasy Plant Mini Kit (Qiagen) and sequenced (Novogene Ltd., UK) as 150 bp paired-end reads on Novaseq (Illumina) to approximately 50x coverage. The mapping was performed using a custom-made bioinformatic protocol as described (Dvořák Tomaštíková et al., 2023).

### Analysis of splicing effects induced by *hze4-1, hze5-4* and *hze6-1* mutations

The effects of *hze4-1*, *hze5-4* and *hze6-1* mutations on splicing were performed by cDNA analysis. Total RNA was isolated from 7-day-old seedlings using an Rneasy Plant Mini kit (Qiagen) with on-column DNAseI digest according to the manufacturer’s instructions, 2 mg of total RNA was used as a template for cDNA synthesis with the RevertAid First Strand cDNA Synthesis kit (K1621, Thermo Scientific) with oligo(dT)12-18 primer according to the kit instructions. The regions of interest were amplified from cDNA by PCR using primer pairs (Table S2) 9A-2_Cf1 and 9A-2_Cr1 for *hze4-1*, zdc1 and zdc2 for *hze5-4*, and caph2-2F and cap-h2-2R for *hze6-1*, and sequenced by Sanger sequencing (SEQme). The sequences were analyzed in SnapGene software and compared to the Col-0 genomic sequence.

### Multiple sequence alignment and protein modeling

Amino acid sequences homologs to *Arabidopsis thaliana* (O24610) CAP-D3 protein identified through BLAST in various species were obtained from the UniProt database (The UniProt Consortium, 2023, https://www.uniprot.org/, February 2024) and Phytozome 13 (46, https://phytozome-next.jgi.doe.gov/, February 2024). Sequences were aligned in the MEGA software (47, https://www.megasoftware.net/) by MUSCLE algorithm, and the results were visualized and prepared for publication using Jalview ver.2 (35, https://www.jalview.org/) with amino acid conservation highlighted according to CLUSTAL conservation of over 50%. Sequences belonging to organisms modeling evolution steps, include: *Brachypodium distachyon* (A0A0Q3EJ37), *Amborella trichopoda* (W1P3N8), *Ceratopteris richardii* (A0A8T2VIA9), *Selaginella moellendorffii* (D8RS24), *Sphagnum magellanicum* (Sphmag07G023100.1.p), *Marchantia polymorpha* (A0A2R6WF77), *Closterium sp.* NIES-68 (A0A9P3IZM4), *Klebsormidium nitens* (A0A1Y1I4J1), *Chlamydomonas reinhardtii* (A0A2K3DT74), *Drosophila melanogaster* (Q9VMQ4), *Xenopus laevis* (Q6GN08), *Danio rerio* (Q7SZF2), *Mus musculus* (Q6ZQK0) and *Homo sapiens* (P42695).

Prediction of the 3D structure of CAP-D3 was created in Alpha-Fold (Jumper et al., 2021). Changes in the structure as a result of amino acid change caused by mutation in hze5-1 were estimated in PyMOL (https://pymol.org/). Global electrostatic potential surfaces were also calculated in PyMOL with the Adaptive Poisson-Boltzmann Solver Tool.

### Cell death assays

Stratified seeds were grown on vertically positioned plates containing ½ MS medium with 0.6 % (w/v) agar. Five-day-old seedlings were transferred to control liquid ½ MS media or media containing DNA-damaging drugs. After 24 h of treatment, the plants were placed onto microscopic slides in a drop of staining solution containing 10 mg ml^-1^ propidium iodide (P4170; Sigma-Aldrich). After mounting a coverslip, the slides were analyzed using the Zeiss AxioImager Z2 microscope (Zeiss) equipped with a high-performance DSD2 confocal module (Andor) and Plan-APOCHROMAT 10x/0.45 objective. For every genotype, 10-15 plants were analyzed, and a representative root was imaged using Leica confocal microscope TCS SP8 with HC PL APO CS2 20x/0.75 DRY objective equipped by LAS-X software with Lightning module laser scanning confocal microscope (all Leica). The images were converted to grayscale in Adobe Photoshop (Adobe Systems).

### Somatic HR assays

The HR trap lines B11 and 651 were crossed with *cap-g2-2* and *cap-h2-1* and double homozygotes were selected in F2 generation. Sterilized F3 double homozygous seeds were grown on ½ MS media plates with 0 (mock), 2.5 µM zebularine and 2.5 µM ICRF-187. Ten-day-old seedlings were histochemically GUS stained overnight at 37 °C as described (Vladejić et al., 2022). The plants were cleared using 70% ethanol washes and numbers of HR events were analyzed under a stereomicroscope. The experiment was performed in three biological replicates with at least 27 plants per genotype.

### Chromosome spreads, FISH and analysis of mitosis

Inflorescences were dissected and put into the liquid ½ MS media supplemented with 40 µM zebularine or 10 µM ICRF-187. After 24 hours, the inflorescences were fixed in Carnoy’s (60% ethanol, 20% chloroform, 10% glacial acetic acid) solution overnight at room temperature, the fixative was exchanged and the samples were stored at −20 °C until use. For analysis, the open flower buds and siliques were removed and the whorl of closed florets was washed in an hourglass 3× 5 min. in distilled water, then 3× 5 min. in 0.1M citric buffer pH 4.6 (0.1M Citric acid monohydrate (68 6318, Chemapol), 0.1M Sodium citrate dihydrate (1.37042, Lachner), and digested in an (w/v) enzyme mixture of 0.3% pectolyase from *Aspergillus japonicus* (P3026; Sigma-Aldrich), 0.3% cytohelicase from *Helix pomatia* (C8274; Sigma-Aldrich) and 0.3% cellulase Onozuka R-10 (16419; Serva) for 4h at 37 °C. Subsequently, the enzyme mixture was replaced with 1× citric buffer, and the samples were either stored at 4 °C or processed directly. Single flower buds were dissected, chopped in 20 μl of 60% acetic acid on a microscopic slide and the suspension was slowly stirred with a needle on a heating block (50 °C) for about 1 min. The slides were fixed by adding 2× 100 μl of Carnoy’s solution, dried and post-fixed in 4% formaldehyde in 1× PBS for 10 min. The slides were air-dried and stored at - 20 °C until use.

FISH probes specific for *45S rDNA* and centromeric repeat (*pAL*) were amplified from *A. thaliana* Col-0 genomic DNA and directly labeled with biotin-dUTP (cat. no. 11093070910; Roche), digoxigenin-dUTP (cat. no. DIGUTP-RO; Roche) or directly labeled dUTP-Cy5 (cat. no. GEPA55022; Sigma-Aldrich) during PCR. Slide pretreatment, hybridization, post-hybridization washes, and detection were carried out as described (Pecinka et al., 2004, 2010). Biotin-dUTP was detected by goat anti-avidin conjugated with biotin (1:100, Vector Laboratories) and avidin combined with Texas Red (1:1000, Vector Laboratories), digoxigenin-dUTP by mouse anti-digoxigenin (1:250, Roche) and goat anti-mouse conjugated with Alexa Fluor 488 (1:200, Molecular Probes). Chromosomes were counterstained with DAPI 1 μg/ml (Vector Laboratories). Slides were examined with the AxioImager Z2 (Zeiss) microscope equipped with a high-performance DSD2 spinning disc module (Andor) and a Cool Cube 1 camera (Metasystems) using appropriate excitation and emission filters. Image processing was carried out using ISIS software 5.4.7 (Metasystems) and Imaris (Bitplane).

### Mutation accumulation and genome resequencing

The homozygous mutant *cap-d3-2* and corresponding WT (segregated out of the heterozygous mutant) plants were propagated under standard conditions (see section Plant materials) for five generations. The genotype was regularly confirmed by PCR based-genotyping with the primer combination: KP118_cap-d3-2F and + KP119_cap-d3-2R (WT allele) and LB_AP1 + KP119_cap-d3-2R (mutant allele) and mutant hypersensitivity to ICRF treatment.

For sequencing, genomic DNA was isolated from three WT siblings and three *cap-d3-*1 siblings in the fifth self-pollinated generation. Illumina libraries were prepared using NEBNext® Ultra™ II DNA Library Prep Kit (Ipswich, MA, USA) and sequenced as 150 bp paired-end reads, ∼30× genome coverage, on the Illumina NovaSeq X Plus platform (Institute of Applied Biotechnologies, Czech Republic). The raw sequence reads were deposited in the NCBI Sequence Read Archive (BioProject ID: PRJNA1106423). Raw data sets were trimmed for quality using Trim Galore https://github.com/FelixKrueger/TrimGalore, mapped onto the Araport11 reference sequence (Cheng et al., 2017) using Bowtie2 (Langmead and Salzberg, 2012), and subsequently treated with GATK (Van der Auwera and O’Connor, 2020), SAMtools (Li et al., 2009) and BCFtools (Li, 2011). To gain a complete set of WT- and mutant-specific single nucleotide polymorphisms (SNPs), we performed a manual curation of the output files in Microsoft Office Excel 2019 (Washington, USA). To filter out any potential segregating pre-accumulation experiment mutations and mutations induced during library preparation, we accepted only polymorphisms that occurred in at least two individuals of the same genotype and simultaneously not present in the other genotype, had a depth of coverage ≥ 20 reads, and allelic depth ≥ 30 % for the alternative allele in comparison with the reference allele.

### Statistical analysis

Statistical analyses were executed in Minitab (Minitab Inc.). Root length measurements were analyzed with a “one-way” analysis of variance (ANOVA) with post hoc Tukey’s test (*P < 0.05*) for root length analysis and Kruskall-Wallis (*P < 0.05*) for HDR assays.

## AUTHOR CONTRIBUTION

AP, KP and EDT designed the research; KP, JV, ZdC, EDT, MP, JZ, KK and AP performed the research; JV, KP, EDT and AP analyzed data; AP wrote the paper with help of all other co-authors. All authors read and approved the manuscript.

## ACKNOWLEDGMENTS

We thank Helena Tvardíková, Eva Jahnová and Kateřina Šléglová for technical assistance, Zdeňka Bursová for plant care, Dr. Kateřina Holušová for sequencing and Petr Navrátil for IT support. This work was supported by the Czech Science Foundation grant 22-00871S to AP and the project TowArds Next GENeration Crops (reg. no. CZ.02.01.01/00/22_008/0004581) of the ERDF Programme Johannes Amos Comenius. Computational resources were provided by the e-INFRA CZ project (ID:90254), supported by the Ministry of Education, Youth and Sports of the Czech Republic.

## CONFLICT OF INTEREST

The authors declare no conflict of interest.

## SUPPORTING INFORMATION

Additional Supporting Information may be found in the online version of this article.

Figure S1. Mapping of the *hze4-1* and *hze4-2* mutations. (A) The *hze4-1* SNP frequency plot of chromosome 5. The position containing the candidate gene is marked by a light red background. (B) Table of *hze4-1* candidate SNPs in the mapped region. A light red highlights the candidate gene At5g62410 (*SMC2A*). REF – nucleotide in reference sequence; ALT – altered nucleotide; PHRED – quality score of the sequencing; MQ – mapping quality. (C) Identification of the chromosomal region containing *hze4-2* mutation in zebularine-sensitive plants selected from F2 *hze4-2* × WT segregating population. The dots represent the mean frequency of five consecutive high-confidence SNPs. Note that no specific candidate region could be identified and an alternative candidate search approach was applied based on the criteria described in (D). (D) Table of *hze4-2* candidate SNPs. The candidate did not show a clear association with a specific region; therefore, we selected high-quality SNPs (PHRED >200) with frequency (PHREQ >65 %) and moderate or high impact. The mutated position in *SMC2A* is marked in light red.

Figure S2. Mapping of the *hze5-1* mutation. (A) Identification of the chromosomal region containing *hze5-1* mutation in zebularine-sensitive plants selected from F2 *hze5-1* × WT segregating population The dots represent the mean frequency of nine consecutive high-confidence SNPs. The candidate region on chromosome 4 (red dots) is highlighted by a light red background. (B) The *hze5-1* SNP frequency plot of chromosome 4. The position containing the candidate gene is marked by a light red background. (C) Table of *hze5-1* candidate SNPs in the mapped region. By light red is highlighted the candidate gene At4g15890 (*CAP-D3*). REF – nucleotide in reference sequence; ALT – altered nucleotide; PHRED – quality score of the sequencing; MQ – mapping quality.

Figure S3. Mapping of the *hze5-2* mutation. (A) Identification of the chromosomal region containing *hze5-2* mutation in zebularine-sensitive plants selected from F2 *hze5-2* × WT segregating population. The dots represent the mean frequency of eleven consecutive high-confidence SNPs. The candidate region on chromosome 4 (red dots) is highlighted by a light red background. (B) The *hze5-2* SNP frequency plot of chromosome 4. The position containing the candidate gene is marked by a light red background. (C) Table of *hze5-2* candidate SNPs in the mapped region. By light red is highlighted the candidate gene At4g15890 (*CAP-D3*). REF – nucleotide in reference sequence; ALT – altered nucleotide; PHRED – quality score of the sequencing; MQ – mapping quality.

Figure S4. Mapping of the *hze5-3* mutation. (A) Identification of the chromosomal region containing *hze5-3* mutation in zebularine-sensitive plants selected from F2 *hze5-3* × WT segregating population. The dots represent the mean frequency of eleven consecutive high-confidence SNPs. The candidate region on chromosome 4 (red dots) is highlighted by a light red background. (B) The *hze5-3* SNP frequency plot of chromosome 4. The position containing the candidate gene is marked by a light red background. (C) Table of *hze5-3* candidate SNPs in the mapped region. By light red is highlighted the candidate gene At4g15890 (*CAP-D3*). REF – nucleotide in reference sequence; ALT – altered nucleotide; PHRED – quality score of the sequencing; MQ – mapping quality.

Figure S5. Mapping of the *hze5-4* mutation. (A) Identification of the chromosomal region containing *hze5-4* mutation in zebularine-sensitive plants selected from F2 *hze5-4* × WT segregating population. The dots represent the mean frequency of eleven consecutive high-confidence SNPs. The candidate region on chromosome 4 (red dots) is highlighted by a light red background. (B) The *hze5-4* SNP frequency plot of chromosome 4. The position containing the candidate gene is marked by a light red background. (C) Table of *hze5-4* candidate SNPs in the mapped region. By light red is highlighted the candidate gene At4g15890 (*CAP-D3*). REF – nucleotide in reference sequence; ALT – altered nucleotide; PHRED – quality score of the sequencing; MQ – mapping quality.

Figure S6. Mapping of the *hze6-1* mutation. (A) Identification of the chromosomal region containing *hze6-1* mutation in zebularine-sensitive plants selected from F2 *hze6-1* × WT segregating population. The dots represent the mean frequency of eleven consecutive high-confidence SNPs. The candidate region on chromosome 3 (red dots) is highlighted by a light red background. (B) The *hze6-1* SNP frequency plot of chromosome 3. The position containing the candidate gene is marked by a light red background. (C) Table of *hze6-1* candidate SNPs in the mapped region. By light red is highlighted the candidate gene At3g16730 (*CAP-H2*). REF – nucleotide in reference sequence; ALT – altered nucleotide; PHRED – quality score of the sequencing; MQ – mapping quality.

Figure S7. Mapping of the *hze6-2* mutation. (A) Identification of the chromosomal region containing *hze6-2* mutation in zebularine-sensitive plants selected from F2 *hze6-2* × WT segregating population. The dots represent the mean frequency of eleven consecutive high-confidence SNPs. The candidate region on chromosome 3 (red dots) is highlighted by a light red background. (B) The *hze6-2* SNP frequency plot of chromosome 3. The position containing the candidate gene is marked by a light red background. (C) Table of *hze6-2* candidate SNPs in the mapped region. By light red is highlighted the candidate gene At3g16730 (*CAP-H2*). REF – nucleotide in reference sequence; ALT – altered nucleotide; PHRED – quality score of the sequencing; MQ – mapping quality.

Figure S8. Condensin II mutants DNA damage sensitivity assays. (A, C, E) Representative seven-day-old plants grown on media without (CTRL), with 50 nM bleomycin (BLE), 10 μM mitomycin C (MMC), 6.5 μM zeocin (ZEO), or 20 nM camptothecin (CPT). Scale bar, 1 cm. (B, D, F) Relative root length of seven days old BLE- and MMC-treated plants. Error bars are the standard deviation of the means of three biological replicates (14-25 plants per replicate). Genotypes marked with the same letter did not differ (P < 0.05) in Tukey’s test.

Figure S9. Condensin II function is independent of SOG1 and SMC5/6 pathways. (A) Relative root length of seven days old ICRF-187 (ICRF)-treated Condensin II and SOG1 double mutant plants. Error bars are the standard deviation of the means of three biological replicates (14-25 plants per replicate). Genotypes marked with the same letter did not differ (P < 0.05) in Tukey’s test. The images of representative plants are provided in Figure 5a. (B) Relative root length of Condensin II and SMC6B double mutant plants. The setup and evaluation were done as described in (A). The images of representative plants are provided in Figure 5d.

Figure S10. Homology-based repair (HDR) in Condensin II mutants.

(A) Frequency of single-strand annealing (SSA) events at the reporter loci B11 and 651 in wild type (WT) and Condensin II mutants under control (CTRL) and 2.5 μM zebularine (ZEB) conditions. Individual dots show the number of SSA events per plant. In total 60 – 105 plants per group were evaluated. The boxplots’ hinges are in the 1st and 3rd quartile, with a marked median. Whisker marks show the lowest or highest value within the 1.5 interquartile range below or above hinges. Statistical significance was tested with the Kruskall-Wallis H-test with *post hoc* Conover-Iman test of multiple comparisons using rank sums with Benjamini-Hochberg procedure (P < ½ α, α = 0.05) in R 4.2.1. *** P < 0.001, NS – not significant.

(B) Frequency of SSA at the reporter loci B11 and 651 after 2.5 μM ICRF-187 (ICRF) treatment. The parameters were as in (A). In total 60 - 105 plants per time point were evaluated.

Table S1. Mutation accumulation experiment. Single nucleotide polymorphisms were identified in the three siblings (R1, R2, R3) of wild-type (WT) and cap-d3-2 mutant (MUT).

Table S2. Primers used in this study.

Table S3. P values of the Chi-squared tests were used to assess the statistical relevance of data presented in Figure 6B.

Table S4. P values of the Chi-squared tests were used to assess the statistical relevance of data presented in Figure 6C.

Table S5. P values of the Chi-squared tests were used to assess the statistical relevance of data presented in Figure 6D.

## Notes

### Competing Interest Statement

The authors have declared no competing interest.

## REFERENCES

Arabidopsis Genome Initiative (2000). Analysis of the genome sequence of the flowering plant Arabidopsis thaliana. Nature 408: 796–815.

Van der Auwera, G. and O’Connor, B.D. (2020). Genomics in the Cloud: Using Docker, GATK, and WDL in Terra 1st Edition 1st Editio. (O’Reilly Media, Inc.).

Baubec, T., Pecinka, A., Rozhon, W., and Mittelsten Scheid, O. (2009). Effective, homogeneous and transient interference with cytosine methylation in plant genomic DNA by zebularine. Plant J. 57: 542–554.

Baxter, J. (2015). “Breaking Up Is Hard to Do”: The Formation and Resolution of Sister Chromatid Intertwines. J. Mol. Biol. 427: 590–607.

Bourbousse, C., Vegesna, N., and Law, J.A. (2018). SOG1 activator and MYB3R repressors regulate a complex DNA damage network in Arabidopsis. Proc. Natl. Acad. Sci. U. S. A. 115: E12453–E12462.

Cheng, C.-Y., Krishnakumar, V., Chan, A.P., Thibaud-Nissen, F., Schobel, S., and Town, C.D. (2017). Araport11: a complete reannotation of the Arabidopsis thaliana reference genome. Plant J. 89: 789–804.

Classen, S., Olland, S., and Berger, J.M. (2003). Structure of the topoisomerase II ATPase region and its mechanism of inhibition by the chemotherapeutic agent ICRF-187. Proc. Natl. Acad. Sci. 100: 10629–10634.

Díaz, M. and Pecinka, A. (2018). Scaffolding for repair: Understanding molecular functions of the SMC5/6 complex. Genes (Basel). 9: 36.

Dixon, J.R., Gorkin, D.U., and Ren, B. (2016). Chromatin Domains: The Unit of Chromosome Organization. Mol. Cell 62: 668–680.

Dvořák Tomaštíková, E. and Pecinka, A. (2025). A Practical Approach to High-Throughput and Accurate Mapping-by-Sequencing in Arabidopsis BT - Methods for Plant Nucleus and Chromatin Studies: Methods and Protocols. In Methods for Plant Nucleus and Chromatin Studies. Methods in Molecular Biology, vol 2873., C. Baroux and C. Tatout, eds (Springer US: New York, NY), pp. 53–70.

Dvořák Tomaštíková, E. and Pecinka, A. (2024). Cytidine analogs in plant epigenetic research and beyond. J. Exp. Bot.: erae522.

Dvořák Tomaštíková, E., Prochazkova, K., Yang, F., Jemelkova, J., Finke, A., Dorn, A., Said, M., Puchta, H., and Pecinka, A. (2023). SMC5/6 complex-mediated SUMOylation stimulates DNA–protein cross-link repair in Arabidopsis. Plant Cell 35: 1532–1547.

Dvořák Tomaštíková, E., Vaculíková, J., Štenclová, V., Kaduchová, K., Pobořilová, Z., Paleček, J.J., and Pecinka, A. (2024). The interplay of homology-directed repair pathways in the repair of zebularine-induced DNA–protein crosslinks in Arabidopsis. Plant J. n/a.

Fransz, P., Soppe, W., and Schubert, I. (2003). Heterochromatin in interphase nuclei of Arabidopsis thaliana. Chromosom. Res. 11: 227–240.

Fraser, J., Williamson, I., Bickmore, W.A., and Dostie, J. (2015). An Overview of Genome Organization and How We Got There: from FISH to Hi-C. Microbiol. Mol. Biol. Rev. 79: 347–372.

Fujimoto, S., Yonemura, M., Matsunaga, S., Nakagawa, T., Uchiyama, S., and Fukui, K. (2005). Characterization and dynamic analysis of Arabidopsis condensin subunits, AtCAP-H and AtCAP-H2. Planta 222: 293–300.

Ganji, M., Shaltiel, I.A., Bisht, S., Kim, E., Kalichava, A., Haering, C.H., and Dekker, C. (2018). Real-time imaging of DNA loop extrusion by condensin. Science (80-.). 360: 102–105.

Goodstein, D.M., Shu, S., Howson, R., Neupane, R., Hayes, R.D., Fazo, J., Mitros, T., Dirks, W., Hellsten, U., Putnam, N., and Rokhsar, D.S. (2012). Phytozome: a comparative platform for green plant genomics. Nucleic Acids Res. 40: D1178–D1186.

Haering, C.H. and Gruber, S. (2016). SnapShot: SMC protein complexes part i. Cell 164: 326–326.e1.

Hirano, T. (2012). Condensins: Universal organizers of chromosomes with diverse functions. Genes Dev. 26: 1659–1678.

Hu, Z., Cools, T., and De Veylder, L. (2016). Mechanisms used by plants to cope with DNA damage. Annu. Rev. Plant Biol. 67: 439–462.

Jeppsson, K., Carlborg, K.K., Nakato, R., Berta, D.G., Lilienthal, I., Kanno, T., Lindqvist, A., Brink, M.C., Dantuma, N.P., Katou, Y., Shirahige, K., and Sjögren, C. (2014). The Chromosomal Association of the Smc5/6 Complex Depends on Cohesion and Predicts the Level of Sister Chromatid Entanglement. PLoS Genet. 10.

Jumper, J. et al. (2021). Highly accurate protein structure prediction with AlphaFold. Nature 596: 583–589.

Langmead, B. and Salzberg, S.L. (2012). Fast gapped-read alignment with Bowtie 2. Nat. Methods 9: 357–359.

Li, H. (2011). A statistical framework for SNP calling, mutation discovery, association mapping and population genetical parameter estimation from sequencing data. Bioinformatics 27: 2987–2993.

Li, H., Handsaker, B., Wysoker, A., Fennell, T., Ruan, J., Homer, N., Marth, G., Abecasis, G., Durbin, R., and Subgroup, 1000 Genome Project Data Processing (2009). The sequence alignment/map format and SAMtools. Bioinformatics 25: 2078– 2079.

Liu, C.H., Finke, A., Díaz, M., Rozhon, W., Poppenberger, B., Baubec, T., and Pecinka, A. (2015). Repair of DNA damage induced by the cytidine analog zebularine requires ATR and ATM in arabidopsis. Plant Cell 27: 1788–1800.

Liu, C.M., McElver, J., Tzafrir, I., Joosen, R., Wittich, P., Patton, D., Van Lammeren, A.A.M., and Meinke, D. (2002). Condensin and cohesin knockouts in Arabidopsis exhibit a titan seed phenotype. Plant J. 29: 405–415.

Molinier, J., Ries, G., Bonhoeffer, S., and Hohn, B. (2004). Interchromatid and Interhomolog Recombination in Arabidopsis thaliana. Plant Cell 16: 342–352.

Municio, C. et al. (2021). The Arabidopsis condensin CAP-D subunits arrange interphase chromatin. New Phytol. n/a.

Nasmyth, K. and Haering, C.H. (2009). Cohesin: Its Roles and Mechanisms. Annu. Rev. Genet. 43: 525–558.

Nitiss, J.L. (2009). Targeting DNA topoisomerase II in cancer chemotherapy. Nat. Rev. Cancer 9: 338–350.

Nowicka, A. et al. (2020). Comparative analysis of epigenetic inhibitors reveals different degrees of interference with transcriptional gene silencing and induction of DNA damage. Plant J 102: 68–84.

Ogita, N. et al. (2018). Identifying the target genes of SUPPRESSOR OF GAMMA RESPONSE 1, a master transcription factor controlling DNA damage response in Arabidopsis. Plant J. 94: 439–453.

Ossowski, S. et al. (2010). The rate and molecular spectrum of spontaneous mutations in Arabidopsis thaliana Supporting Online Material.

Pecinka, A., Dinh, H.Q., Baubec, T., Rosa, M., Lettner, N., and Mittelsten Scheid, O. (2010). Epigenetic regulation of repetitive elements is attenuated by prolonged heat stress in Arabidopsis. Plant Cell 22: 3118–3129.

Pecinka, A., Rosa, M., Schikora, A., Berlinger, M., Hirt, H., Luschnig, C., and Scheid, O.M. (2009). Transgenerational stress memory is not a general response in Arabidopsis. PLoS One 4.

Pecinka, A., Schubert, V., Meister, A., Kreth, G., Klatte, M., Lysak, M.A., Fuchs, J., and Schubert, I. (2004). Chromosome territory arrangement and homologous pairing in nuclei of Arabidopsis thaliana are predominantly random except for NOR-bearing chromosomes. Chromosoma 113: 258–269.

Prochazkova, K., Finke, A., Tomaštíková, E.D., Filo, J., Bente, H., Dvořák, P., Ovečka, M., Šamaj, J., and Pecinka, A. (2021). Zebularine induces enzymatic DNA–protein crosslinks in 45S rDNA heterochromatin of Arabidopsis nuclei. Nucleic Acids Res.: gkab1218.

Puchta, H., Swoboda, P., and Hohn, B. (1995). Induction of intrachromosomal homologous recombination in whole plants. Plant J. 7: 203–210.

Roy, S., Adhikary, H., and D’Amours, D. (2024). The SMC5/6 complex: folding chromosomes back into shape when genomes take a break. Nucleic Acids Res. 52: 2112–2129.

Sakamoto, T. et al. (2022). Two-step regulation of centromere distribution by condensin II and the nuclear envelope proteins. Nat. Plants 8: 940–953.

Sakamoto, T., Inui, Y.T., Uraguchi, S., Yoshizumi, T., Matsunaga, S., Mastui, M., Umeda, M., Fukui, K., and Fujiwara, T. (2011). Condensin II alleviates DNA damage and is essential for tolerance of boron overload stress in arabidopsis. Plant Cell 23: 3533– 3546.

Sakamoto, T., Sugiyama, T., Yamashita, T., and Matsunaga, S. (2019). Plant condensin II is required for the correct spatial relationship between centromeres and rDNA arrays. Nucleus 10: 116–125.

Schubert, I. and Oud, J.L. (1997). There Is an Upper Limit of Chromosome Size for Normal Development of an Organism. Cell 88: 515–520.

Schubert, V. (2009). SMC proteins and their multiple functions in higher plants. Cytogenet. Genome Res. 124: 202–214.

Sen, N., Leonard, J., Torres, R., Garcia-Luis, J., Palou-Marin, G., and Aragón, L. (2016). Physical Proximity of Sister Chromatids Promotes Top2-Dependent Intertwining. Mol. Cell 64: 134–147.

Siddiqui, N.U., Rusyniak, S., Hasenkampf, C.A., and Riggs, C.D. (2006). Disruption of the Arabidopsis SMC4 gene, AtCAP-C, compromises gametogenesis and embryogenesis. Planta 223: 990–997.

Siddiqui, N.U., Stronghill, P.E., Dengler, R.E., Hasenkampf, C.A., and Riggs, C.D. (2003). Mutations in Arabidopsis condensin genes disrupt embryogenesis, meristem organization and segregation of homologous chromosomes during meiosis. Development 130: 3283–3295.

Smith, S.J., Osman, K., and Franklin, F.C.H. (2014). The condensin complexes play distinct roles to ensure normal chromosome morphogenesis during meiotic division in Arabidopsis. Plant J. 80: 255–268.

Stingele, J., Bellelli, R., and Boulton, S.J. (2017). Mechanisms of DNA-protein crosslink repair. Nat. Rev. Mol. Cell Biol. 18: 563–573.

Tamura, K., Stecher, G., and Kumar, S. (2021). MEGA11: Molecular Evolutionary Genetics Analysis Version 11. Mol. Biol. Evol. 38: 3022–3027.

Uhlmann, F. (2016). SMC complexes: From DNA to chromosomes. Nat. Rev. Mol. Cell Biol. 17: 399–412.

Vladejić, J., Yang, F., Dvořák Tomaštíková, E., Doležel, J., Palecek, J.J., and Pecinka, A. (2022). Analysis of BRCT5 domain-containing proteins reveals a new component of DNA damage repair in Arabidopsis. Front. Plant Sci. 13.

Watanabe, K., Pacher, M., Dukowic, S., Schubert, V., Puchta, H., and Schubert, I. (2009). The STRUCTURAL MAINTENANCE of CHROMOSOMES 5/6 complex promotes sister chromatid alignment and homologous recombination after DNA damage in Arabidopsis thaliana. Plant Cell 21: 2688–2699.

Waterhouse, A.M., Procter, J.B., Martin, D.M.A., Clamp, M., and Barton, G.J. (2009). Jalview Version 2—a multiple sequence alignment editor and analysis workbench. Bioinformatics 25: 1189–1191.

Willing, E.M., Piofczyk, T., Albert, A., Winkler, J.B., Schneeberger, K., and Pecinka, A. (2016). UVR2 ensures transgenerational genome stability under simulated natural UV-B in Arabidopsis thaliana. Nat. Commun. 7: 1–9.

Yoshiyama, K., Conklin, P.A., Huefner, N.D., and Britt, A.B. (2009). Suppressor of gamma response 1 (SOG1) encodes a putative transcription factor governing multiple responses to DNA damage. Proc. Natl. Acad. Sci. U. S. A. 106: 12843–12848.

