## Supplemental data for "Condensin II mediates efficient chromatid resolution and resistance to genotoxic stress in Arabidopsis"

SUPPLEMENTARY MATERIALS

A

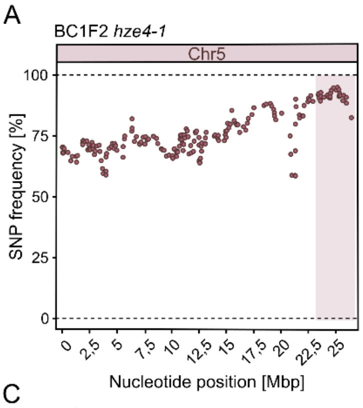


B

| Chr | Nucleotide position | Ref | Alt | Freq | Phred | MQ | Effect | Impact | AA change | Gene | Gene name |
|---|---|---|---|---|---|---|---|---|---|---|---|
| Chr5 | 24619197 | G | A | 88,06 | 219 | 41 | synonymous_variant | LOW | R804 | AT5G61190 | AT5G61190 |
| Chr5 | 24805638 | G | A | 96,36 | 228 | 39 | missense_variant | MODERATE | T531I | AT5G61730 | ABCA9 |
| Chr5 | 25061626 | G | A | 92,00 | 228 | 41 | splice_donor_variant intron_variant | HIGH | | AT5G62410 | SMC2 |
| Chr5 | 25164917 | G | A | 97,87 | 228 | 42 | missense_variant | MODERATE | G944D | AT5G62670 | AHA11 |
| Chr5 | 25296966 | G | A | 94,83 | 228 | 41 | upstream_gene_variant | MODIFIER | | AT5G63060 | CPSFL1 |
| Chr5 | 25307165 | G | A | 92,59 | 228 | 41 | upstream_gene_variant | MODIFIER | | AT5G63080 | JMJ20 |
| Chr5 | 25339120 | G | A | 91,30 | 223 | 42 | missense_variant | MODERATE | L230F | AT5G63170 | GDSL-like |
| Chr5 | 25370401 | G | A | 98,08 | 228 | 42 | missense_variant | MODERATE | S165L | AT5G63290 | CPO3 |

C

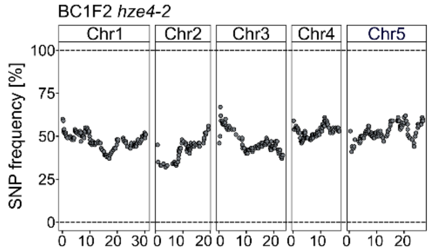


D

| Chr | Nucleotide position | Ref | Alt | Freq | Phred | MQ | Effect | Impact | AA change | Gene | Gene name |
|---|---|---|---|---|---|---|---|---|---|---|---|
| Chr4 | 11451846 | C | T | 71,43 | 222 | 42 | missense_variant | MODERATE | A153V | AT4G21530 | APC4 |
| Chr5 | 25057262 | C | T | 71,43 | 222 | 42 | stop_gained | HIGH | Q286* | AT5G62410 | SMC2-1 |
| Chr2 | 19303204 | C | T | 65,63 | 222 | 42 | missense_variant | MODERATE | A76T | AT2G46980 | ASY3 |
| Chr5 | 19260162 | C | T | 65,63 | 222 | 41 | missense_variant | MODERATE | R762W | AT5G47480 | MAG5 |
| Chr4 | 14601939 | C | T | 65,38 | 222 | 41 | missense_variant splice_region_variant | MODERATE | G64E | AT4G29850 | AT4G29850 |
| Chr5 | 3874775 | G | A | 65,38 | 222 | 41 | missense_variant | MODERATE | A522V | AT5G12000 | AT5G12000 |

**Figure S1.** Mapping of the *hze4-1* and *hze4-2* mutations.

(**A**) The *hze4-1* SNP frequency plot of chromosome 5. The position containing the candidate gene is marked by a light red background.

(**B**) Table of *hze4-1* candidate SNPs in the mapped region. A light red highlights the candidate gene At5g62410 (*SMC2A*). REF – nucleotide in reference sequence; ALT – altered nucleotide; PHRED – quality score of the sequencing; MQ – mapping quality.

(**C**) Identification of the chromosomal region containing *hze4-2* mutation in zebularine-sensitive plants selected from F2 *hze4-2* × WT segregating population. The dots represent the mean frequency of five consecutive high-confidence SNPs. Note that no specific candidate region could be identified and an alternative candidate search approach was applied based on the criteria described in (D).

(**D**) Table of *hze4-2* candidate SNPs. The candidate did not show a clear association with a specific region; therefore, we selected high-quality SNPs (PHRED >200) with frequency (PHREQ >65 %) and moderate or high impact. The mutated position in *SMC2A* is marked in light red.

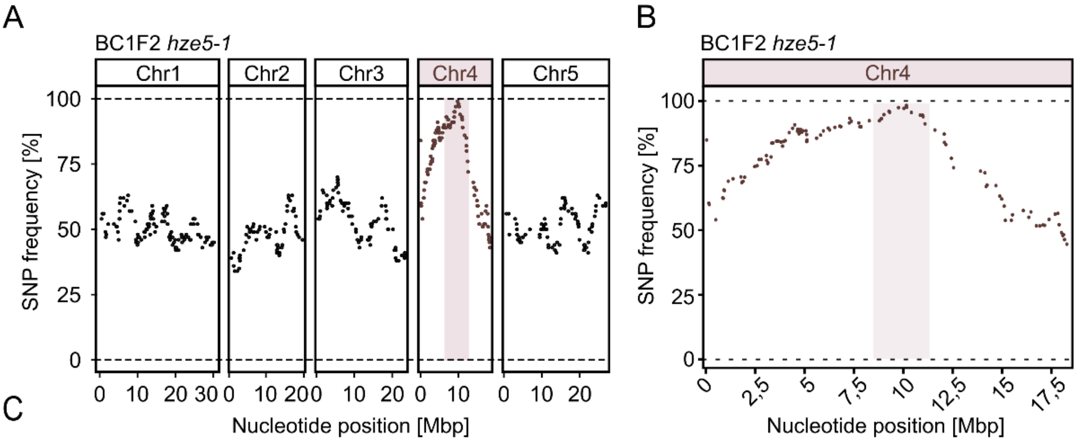


C

BC1F2 *hze5-1*

| Chr | Nucleotide position | Ref | Alt | Freq | Phred | MQ | Effect | Impact | AA change | Gene | Gene name |
|---|---|---|---|---|---|---|---|---|---|---|---|
| Chr4 | 8790860 | G | A | 100 | 228 | 42 | downstream_gene_variant | MODIFIER | | AT4G15380 | CYP705A4 |
| Chr4 | 8885899 | G | A | 100 | 228 | 42 | missense_variant | MODERATE | D375N | AT4G15560 | DXS |
| Chr4 | 8895553 | G | A | 100 | 228 | 42 | splice_region_variant&intron_variant | LOW | | AT4G15570 | MAA3 |
| Chr4 | 9019655 | G | A | 100 | 228 | 42 | misense_variant | MODERATE | G631D | AT4G15890 | CAP-D3 |
| Chr4 | 9267402 | G | A | 100 | 228 | 42 | 5_prime_UTR_variant | MODIFIER | | AT4G16430 | BHLH3 |
| Chr4 | 9314157 | G | A | 95,83 | 228 | 42 | missense_variant | MODERATE | R146K | AT4G16535 | AT4G16535 |
| Chr4 | 10029865 | G | A | 89,47 | 228 | 42 | 5_prime_UTR_premature_start_codon_gain_variant | LOW | | AT4G18060 | SH3P3 |
| Chr4 | 10408554 | G | A | 77,59 | 221 | 42 | missense_variant | MODERATE | P10L | AT4G19003 | VPS25 |
| Chr4 | 10856800 | G | A | 88,57 | 223 | 42 | synonymous_variant | LOW | F465F | AT4G20060 | EMB1895 |

**Figure S2.** Mapping of the *hze5-1* mutation.

(**A**) Identification of the chromosomal region containing *hze5-1* mutation in zebularine-sensitive plants selected from F2 *hze5-1* × WT segregating population The dots represent the mean frequency of nine consecutive high-confidence SNPs. The candidate region on chromosome 4 (red dots) is highlighted by a light red background.

(**B**) The *hze5-1* SNP frequency plot of chromosome 4. The position containing the candidate gene is marked by a light red background.

(**C**) Table of *hze5-1* candidate SNPs in the mapped region. By light red is highlighted the candidate gene At4g15890 (*CAP-D3*). REF – nucleotide in reference sequence; ALT – altered nucleotide; PHRED – quality score of the sequencing; MQ – mapping quality.

A B

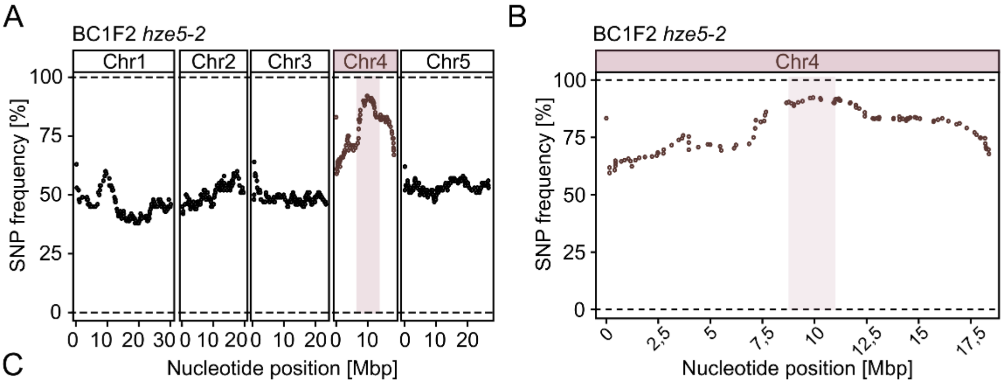


C

BC1F2 *hze5-2*

| Chr | Nucleotide position | Ref | Alt | Freq | Phred | MQ | Effect | Impact | AA change | Gene | Gene name |
|---|---|---|---|---|---|---|---|---|---|---|---|
| Chr4 | 7069795 | C | T | 94,12 | 228 | 42 | intron_variant | MODIFIER | | AT4G11730 | AT4G11730 |
| Chr4 | 7148460 | C | T | 87,50 | 219 | 42 | missense_variant | MODERATE | T34I | AT4G11890 | AT4G11890 |
| Chr4 | 7182943 | C | T | 84,85 | 220 | 42 | missense_variant | MODERATE | A442V | AT4G11970 | AT4G11970 |
| Chr4 | 7584720 | C | T | 85,71 | 219 | 42 | 3_prime_UTR_variant | MODIFIER | | AT4G12960 | *GILT* |
| Chr4 | 7621169 | C | T | 94,74 | 228 | 42 | upstream_gene_variant | MODIFIER | | AT4G13060 | AT4G13060 |
| Chr4 | 7671592 | C | T | 96,15 | 228 | 42 | synonymous_variant | LOW | L57L | AT4G13210 | AT4G13210 |
| Chr4 | 8650412 | C | T | 100 | 228 | 42 | downstream_gene_variant | MODIFIER | | AT4G15165 | *PAC2* |
| Chr4 | 8770003 | C | T | 97,50 | 228 | 42 | upstream_gene_variant | MODIFIER | | AT4G15360 | *CYP705A3* |
| Chr4 | 8893157 | C | T | 87,23 | 219 | 42 | splice_region_variant intron_variant | LOW | | AT4G15570 | *MAA3* |
| Chr4 | 9021857 | C | T | 100 | 228 | 42 | stop_gained | HIGH | Q1045* | AT4G15890 | *CAP-D3* |
| Chr4 | 9214480 | C | T | 97,30 | 228 | 42 | splice_region_variant intron_variant | LOW | | AT4G16280 | *FCA* |
| Chr4 | 9790227 | C | T | 90,24 | 220 | 42 | stop_gained | HIGH | W162* | AT4G17580 | AT4G17580 |
| Chr4 | 10280945 | C | T | 91,23 | 228 | 42 | upstream_gene_variant | MODIFIER | | AT4G18680 | AT4G18680 |
| Chr4 | 10305062 | C | T | 90,63 | 227 | 42 | synonymous_variant | LOW | F71F | AT4G18750 | *DOT4* |
| Chr4 | 10916307 | C | T | 84,62 | 220 | 42 | intron_variant | MODIFIER | | AT4G20210 | *TPS08* |
| Chr4 | 10952209 | C | T | 96,88 | 228 | 42 | synonymous_variant | LOW | V796V | AT4G20270 | *BAM3* |
| Chr4 | 10959264 | C | T | 87,04 | 219 | 42 | downstream_gene_variant | MODIFIER | | AT4G20300 | AT4G20300 |
| Chr4 | 11020661 | C | T | 100 | 228 | 42 | intron_variant | MODIFIER | | AT4G20430 | *SBT2.2* |
| Chr4 | 11135160 | C | T | 100 | 228 | 41 | splice_region_variant intron_variant | LOW | | AT4G20790 | AT4G20790 |
| Chr4 | 11166247 | C | T | 92,11 | 228 | 42 | synonymous_variant | LOW | R653R | AT4G20850 | *TPP2* |
| Chr4 | 11183927 | C | T | 80,95 | 220 | 42 | 3_prime_UTR_variant | MODIFIER | | AT4G20890 | *TUBB9* |
| Chr4 | 11240266 | C | T | 90 | 218 | 42 | upstream_gene_variant | MODIFIER | | AT4G21060 | AT4G21060 |

**Figure S3.** Mapping of the *hze5-2* mutation.
(**A**) Identification of the chromosomal region containing *hze5-2* mutation in zebularine-sensitive plants selected from F2 *hze5-2* × WT segregating population. The dots represent the mean frequency of eleven consecutive high-confidence SNPs. The candidate region on chromosome 4 (red dots) is highlighted by a light red background.
(**B**) The *hze5-2* SNP frequency plot of chromosome 4. The position containing the candidate gene is marked by a light red background.
(**C**) Table of *hze5-2* candidate SNPs in the mapped region. By light red is highlighted the candidate gene At4g15890 (*CAP-D3*). REF – nucleotide in reference sequence; ALT – altered nucleotide; PHRED – quality score of the sequencing; MQ – mapping quality.

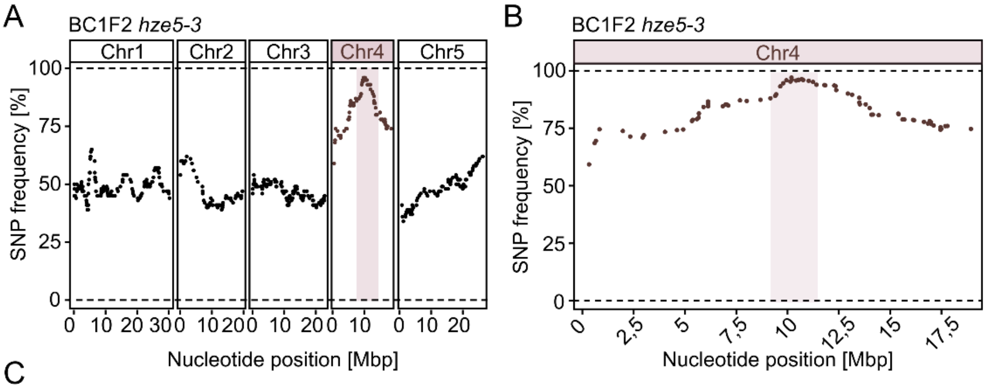


C

BC1F2 *hze5-3*

| Chr | Nucleotide position | Ref | Alt | Freq | Phred | MQ | Effect | Impact | AA change | Gene | Gene name |
|---|---|---|---|---|---|---|---|---|---|---|---|
| Chr4 | 8304249 | G | A | 96,55 | 228 | 42 | upstream_gene_variant | MODIFIER | | AT4G14420 | AT4G14420 |
| Chr4 | 8790044 | G | A | 97,30 | 228 | 42 | missense_variant | MODERATE | V403M | AT4G15380 | *CYP705A4* |
| Chr4 | 9019731 | G | A | 96,00 | 228 | 42 | stop_gained | HIGH | W656* | AT4G15890 | *CAP-D3* |
| Chr4 | 9069963 | C | NA | 96,43 | 228 | 42 | NA | NA | NA | NA | NA |
| Chr4 | 9266796 | G | A | 100,00 | 228 | 42 | upstream_gene_variant | MODIFIER | | AT4G16430 | *BHLH3* |
| Chr4 | 9411570 | G | A | 96,43 | 228 | 42 | upstream_gene_variant | MODIFIER | | AT4G16745 | AT4G16745 |
| Chr4 | 9567748 | G | A | 100,00 | 228 | 42 | intron_variant | MODIFIER | | AT4G17000 | AT4G17000 |
| Chr4 | 9764368 | G | A | 93,75 | 228 | 42 | downstream_gene_variant | MODIFIER | | AT4G17505 | AT4G17505 |
| Chr4 | 9815443 | G | A | 96,77 | 228 | 42 | upstream_gene_variant | MODIFIER | | AT4G17612 | AT4G17612 |
| Chr4 | 9829807 | G | A | 89,19 | 223 | 42 | downstream_gene_variant | MODIFIER | | AT4G17650 | AT4G17650 |
| Chr4 | 10043606 | G | A | 90,63 | 221 | 42 | missense_variant | MODERATE | P746S | AT4G18130 | *PHYE* |
| Chr4 | 10158078 | G | A | 100,00 | 228 | 42 | missense_variant | MODERATE | G41D | AT4G18380 | AT4G18380 |
| Chr4 | 10172594 | C | NA | 96,67 | 228 | 32 | NA | NA | NA | NA | NA |
| Chr4 | 10314793 | G | A | 100,00 | 228 | 41 | missense_variant | MODERATE | P439L | AT4G18780 | *CESA8* |
| Chr4 | 10383806 | G | A | 91,89 | 228 | 42 | intron_variant | MODIFIER | | AT4G18960 | *AG* |
| Chr4 | 10654875 | G | A | 96,30 | 228 | 42 | missense_variant | MODERATE | G547S | AT4G19530 | AT4G19530 |
| Chr4 | 10864757 | G | A | 90,63 | 221 | 42 | 3_prime_UTR_variant | MODIFIER | | AT4G20070 | *AAH* |
| Chr4 | 11032780 | G | A | 86,96 | 219 | 42 | upstream_gene_variant | MODIFIER | | AT4G20460 | AT4G20460 |
| Chr4 | 11418842 | G | C | 92,31 | 228 | 42 | missense_variant | MODERATE | N212K | AT4G21440 | *ATMYB102* |
| Chr4 | 11460946 | G | A | 95,24 | 228 | 42 | missense_variant | MODERATE | G229E | AT4G21540 | *SPHK1* |
| Chr4 | 11901136 | G | A | 88,57 | 219 | 41 | upstream_gene_variant | MODIFIER | | AT4G22600 | *INP1* |
| Chr4 | 11963997 | G | A | 87,80 | 219 | 42 | missense_variant | MODERATE | A296V | AT4G22770 | *AHL2* |
| Chr4 | 11997423 | G | A | 80,77 | 220 | 42 | 5_prime_UTR_variant | MODIFIER | | AT4G22860 | AT4G22860 |
| Chr4 | 12563139 | G | A | 80,00 | 220 | 41 | upstream_gene_variant | MODIFIER | | AT4G24204 | AT4G24204 |
| Chr4 | 12768346 | G | A | 94,59 | 228 | 42 | upstream_gene_variant | MODIFIER | | AT4G24770 | *CP31A* |

**Figure S4.** Mapping of the *hze5-3* mutation.

(**A**) Identification of the chromosomal region containing *hze5-3* mutation in zebularine-sensitive plants selected from F2 *hze5-3* × WT segregating population. The dots represent the mean frequency of eleven consecutive high-confidence SNPs. The candidate region on chromosome 4 (red dots) is highlighted by a light red background.

(**B**) The *hze5-3* SNP frequency plot of chromosome 4. The position containing the candidate gene is marked by a light red background.

(**C**) Table of *hze5-3* candidate SNPs in the mapped region. By light red is highlighted the candidate gene At4g15890 (*CAP-D3*). REF – nucleotide in reference sequence; ALT – altered nucleotide; PHRED – quality score of the sequencing; MQ – mapping quality.

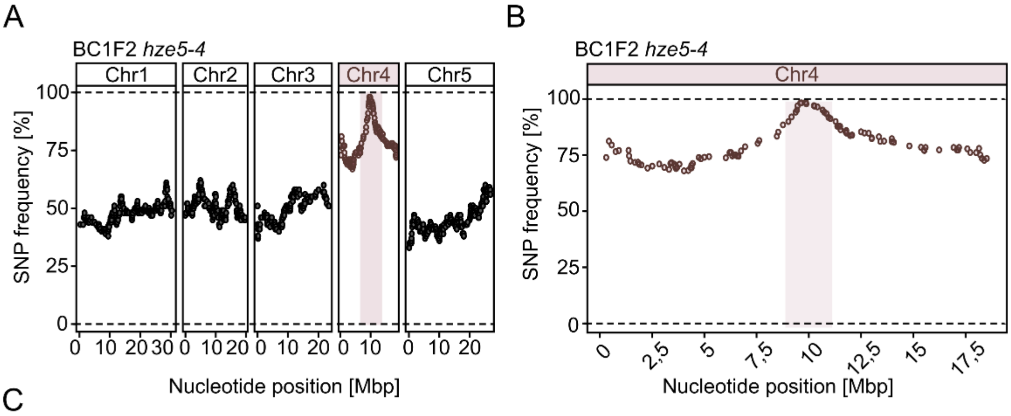


C

BC1F2 *hze5-4*

| Chr | Nucleotide position | Ref | Alt | Freq | Phred | MQ | Effect | Impact | AA change | Gene | Gene name |
|---|---|---|---|---|---|---|---|---|---|---|---|
| Chr4 | 8470620 | G | A | 100 | 228 | 42 | missense_variant | MODERATE | A117T | AT4G14750 | *IQD19* |
| Chr4 | 8536077 | G | A | 100 | 228 | 42 | 5_prime_UTR_variant | MODIFIER | | AT4G14920 | AT4G14920 |
| Chr4 | 8763358 | G | A | 95,24 | 228 | 42 | missense_variant | MODERATE | V136I | AT4G15350 | *CYP705A2* |
| Chr4 | 9021998 | G | A | 100 | 228 | 42 | splice_donor_variant intron_variant | HIGH | | AT4G15890 | *CAP-D3* |
| Chr4 | 9172957 | G | A | 100 | 228 | 42 | 3_prime_UTR_variant | MODIFIER | | AT4G16190 | *RD19C* |
| Chr4 | 9370615 | G | A | 100 | 228 | 42 | upstream_gene_variant | MODIFIER | | AT4G06405 | AT4G06405 |
| Chr4 | 9434159 | G | A | 97,87 | 228 | 42 | 3_prime_UTR_variant | MODIFIER | | AT4G16770 | AT4G16770 |
| Chr4 | 9460188 | G | A | 100 | 228 | 42 | stop_gained | HIGH | Q263* | AT4G16810 | AT4G16810 |
| Chr4 | 9590395 | G | A | 93,75 | 228 | 42 | splice_donor_variant intron_variant | HIGH | | AT4G17050 | *UGLYAH* |
| Chr4 | 9639336 | G | A | 97,5 | 228 | 42 | splice_region_variant synonymous_variant | LOW | K128K | AT4G17150 | AT4G17150 |
| Chr4 | 9858955 | G | A | 100 | 228 | 42 | missense_variant | MODERATE | P84L | AT4G17710 | *HDG4* |
| Chr4 | 9881351 | G | A | 91,43 | 228 | 42 | synonymous_variant | LOW | S35S | AT4G17780 | AT4G17780 |
| Chr4 | 10244436 | C | T | 91,18 | 225 | 42 | synonymous_variant | LOW | P564P | AT4G18600 | *WAVE5* |
| Chr4 | 10276051 | G | A | 92,86 | 228 | 42 | missense_variant | MODERATE | A814V | AT4G18670 | *LRX5* |
| Chr4 | 10412226 | G | A | 96,55 | 228 | 42 | missense_variant | MODERATE | A470V | AT4G19010 | *4CLL6* |
| Chr4 | 10662012 | G | A | 84,37 | 220 | 42 | missense_variant | MODERATE | E37K | AT4G19560 | *CYCT1-2* |
| Chr4 | 10740097 | G | A | 89,47 | 218 | 42 | missense_variant | MODERATE | T69M | AT4G19740 | AT4G19740 |
| Chr4 | 10845562 | G | A | 90,48 | 223 | 42 | intron_variant | MODIFIER | | AT4G20020 | *MORF1* |
| Chr4 | 10943904 | G | A | 81,82 | 220 | 42 | downstream_gene_variant | MODIFIER | | AT4G20260 | *PCAP1* |
| Chr4 | 11033294 | G | NA | 87,88 | 219 | 37 | NA | NA | NA | NA | NA |
| Chr4 | 11281145 | G | A | 83,34 | 220 | 42 | intron_variant | MODIFIER | | AT4G21150 | *RPN2* |

**Figure S5.** Mapping of the *hze5-4* mutation.
(**A**) Identification of the chromosomal region containing *hze5-4* mutation in zebularine-sensitive plants selected from F2 *hze5-4* × WT segregating population. The dots represent the mean frequency of eleven consecutive high-confidence SNPs. The candidate region on chromosome 4 (red dots) is highlighted by a light red background.
(**B**) The *hze5-4* SNP frequency plot of chromosome 4. The position containing the candidate gene is marked by a light red background.
(**C**) Table of *hze5-4* candidate SNPs in the mapped region. By light red is highlighted the candidate gene At4g15890 (*CAP-D3*). REF – nucleotide in reference sequence; ALT – altered nucleotide; PHRED – quality score of the sequencing; MQ – mapping quality.

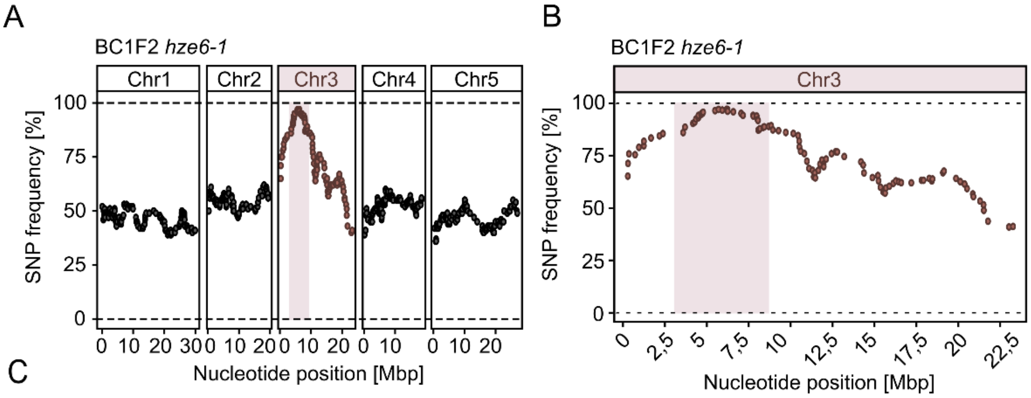


C

BC1F2 *hze6-1*

| Chr | Nucleotide position | Ref | Alt | Freq | Phred | MQ | Effect | Impact | AA change | Gene name | Gene |
|---|---|---|---|---|---|---|---|---|---|---|---|
| Chr3 | 4663121 | C | T | 100 | 228 | 42 | 5_prime_UTR_variant | MODIFIER | | AT3G14070 | *CCX3* |
| Chr3 | 4697491 | C | T | 100 | 228 | 42 | splice_region_variant intron_variant | LOW | | AT3G14170 | AT3G14170 |
| Chr3 | 4745431 | C | T | 93,10 | 228 | 42 | downstream_gene_variant | MODIFIER | | AT3G14250 | AT3G14250 |
| Chr3 | 4778508 | C | T | 100 | 228 | 42 | upstream_gene_variant | MODIFIER | | AT3G14320 | *ATL61* |
| Chr3 | 5491571 | C | T | 100 | 228 | 42 | 3_prime_UTR_variant | MODIFIER | | AT3G16200 | AT3G16200 |
| Chr3 | 5696990 | C | T | 100 | 228 | 41 | splice_donor_variant intron_variant | HIGH | | AT3G16730 | *CAP-H2* |
| Chr3 | 5921337 | C | T | 93,55 | 228 | 42 | synonymous_variant | LOW | R944R | AT3G17340 | AT3G17340 |
| Chr3 | 6182364 | C | T | 96,00 | 228 | 42 | missense_variant | MODERATE | A260V | AT3G18050 | AT3G18050 |
| Chr3 | 6233229 | C | T | 90,00 | 225 | 42 | missense_variant | MODERATE | T335I | AT3G18190 | *CCT4* |
| Chr3 | 6240399 | C | T | 90,24 | 220 | 42 | splice_acceptor_variant intron_variant | HIGH | | AT3G18210 | AT3G18210 |
| Chr3 | 6579639 | C | T | 92,31 | 228 | 41 | intron_variant | MODIFIER | | AT3G19050 | *KIN12D* |

**Figure S6.** Mapping of the *hze6-1* mutation.

(**A**) Identification of the chromosomal region containing *hze6-1* mutation in zebularine-sensitive plants selected from F2 *hze6-1* × WT segregating population. The dots represent the mean frequency of eleven consecutive high-confidence SNPs. The candidate region on chromosome 3 (red dots) is highlighted by a light red background.

(**B**) The *hze6-1* SNP frequency plot of chromosome 3. The position containing the candidate gene is marked by a light red background.

(**C**) Table of *hze6-1* candidate SNPs in the mapped region. By light red is highlighted the candidate gene At3g16730 (*CAP-H2*). REF – nucleotide in reference sequence; ALT – altered nucleotide; PHRED – quality score of the sequencing; MQ – mapping quality.

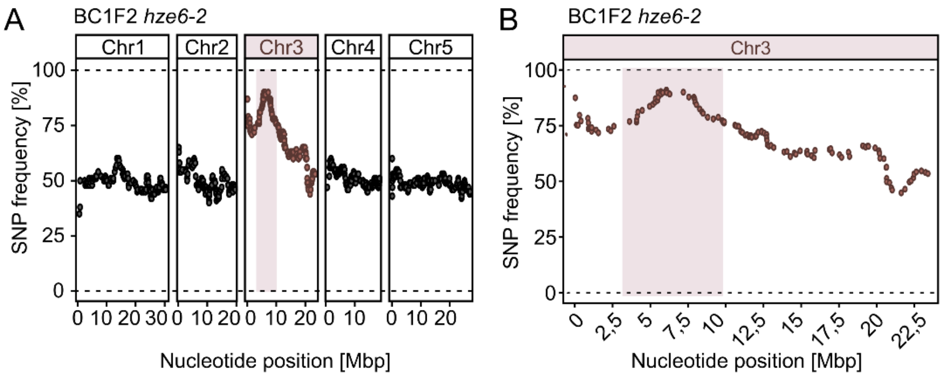


C

BC1F2 *hze6-2*

| Chr | Nucleotide position | Ref | Alt | Freq | Phred | MQ | Effect | Impact | AA change | Gene name | Gene |
|---|---|---|---|---|---|---|---|---|---|---|---|
| Chr3 | 5175064 | C | T | 89,28 | 219 | 42 | missense_variant | MODERATE | A119T | AT3G15356 | *LEC* |
| Chr3 | 5310715 | C | T | 95,65 | 228 | 41 | missense_variant | MODERATE | G35D | AT3G15670 | *LEA29* |
| Chr3 | 5348355 | C | T | 93,34 | 228 | 42 | missense_variant | MODERATE | G121E | AT3G15810 | AT3G15810 |
| Chr3 | 5479888 | C | T | 77,78 | 221 | 42 | intron_variant | MODIFIER | | AT3G16170 | *AAE13* |
| Chr3 | 5521799 | C | T | 93,94 | 228 | 42 | intron_variant | MODIFIER | | AT3G16290 | *FTSHI2* |
| Chr3 | 5523681 | C | T | 95,35 | 228 | 41 | missense_variant | MODERATE | G439R | AT3G16290 | *FTSHI2* |
| Chr3 | 5697055 | C | T | 88,57 | 219 | 41 | stop_gained | HIGH | W359* | AT3G16730 | *CAP-H2* |
| Chr3 | 5698094 | C | T | 80 | 221 | 42 | missense_variant | MODERATE | G103S | AT3G16730 | *CAP-H2* |
| Chr3 | 5699790 | C | T | 93,75 | 228 | 41 | synonymous_variant | LOW | I105I | AT3G16740 | AT3G16740 |
| Chr3 | 5826435 | C | T | 96,43 | 228 | 42 | upstream_gene_variant | MODIFIER | | AT3G17090 | AT3G17090 |
| Chr3 | 6066659 | C | T | 96,55 | 228 | 42 | downstream_gene_variant | MODIFIER | | AT3G17730 | *ANAC057* |
| Chr3 | 6087117 | C | T | 85,71 | 219 | 42 | synonymous_variant | LOW | I34I | AT3G17780 | AT3G17780 |
| Chr3 | 6089534 | C | T | 86,84 | 219 | 42 | 5_prime_UTR_variant | MODIFIER | | AT3G17790 | *PAP17* |
| Chr3 | 6286700 | C | T | 86,21 | 219 | 42 | missense_variant | MODERATE | E149K | AT3G18310 | AT3G18310 |
| Chr3 | 7188871 | G | A | 86,67 | 219 | 42 | missense_variant | MODERATE | P404L | AT3G20580 | *COBL10* |

**Figure S7.** Mapping of the *hze6-2* mutation.

(**A**) Identification of the chromosomal region containing *hze6-2* mutation in zebularine-sensitive plants selected from F2 *hze6-2* × WT segregating population. The dots represent the mean frequency of eleven consecutive high-confidence SNPs. The candidate region on chromosome 3 (red dots) is highlighted by a light red background.

(**B**) The *hze6-2* SNP frequency plot of chromosome 3. The position containing the candidate gene is marked by a light red background.

(**C**) Table of *hze6-2* candidate SNPs in the mapped region. By light red is highlighted the candidate gene At3g16730 (*CAP-H2*). REF – nucleotide in reference sequence; ALT – altered nucleotide; PHRED – quality score of the sequencing; MQ – mapping quality.

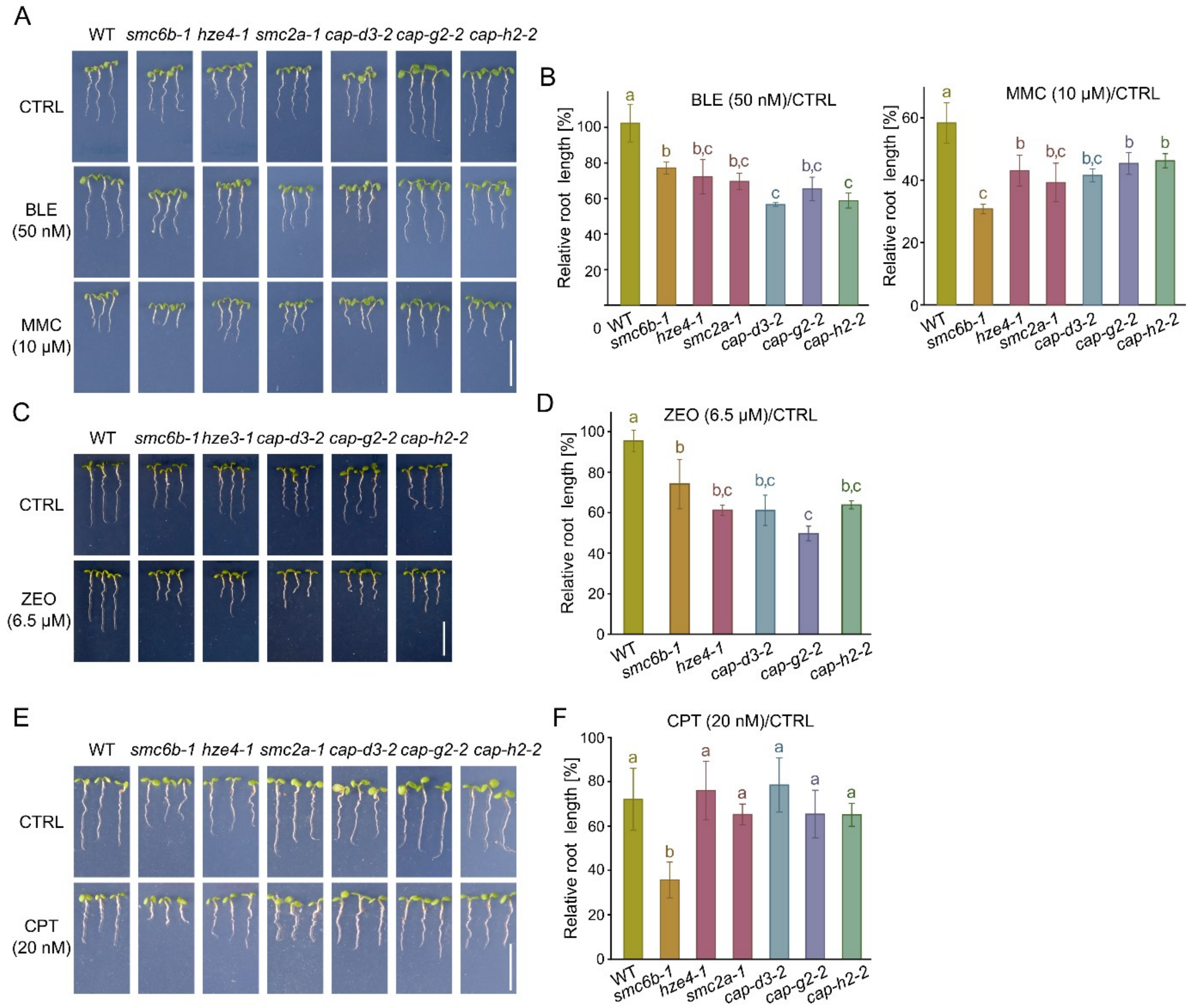


**Figure S8.** Condensin II mutants DNA damage sensitivity assays.
(**A, C, E**) Representative seven-day-old plants grown on media without (CTRL), with 50 nM bleomycin (BLE), 10 µM mitomycin C (MMC), 6.5 µM zeocin (ZEO), or 20 nM camptothecin (CPT). Scale bar, 1 cm.
(**B, D, F**) Relative root length of seven days old BLE- and MMC-treated plants. Error bars are the standard deviation of the means of three biological replicates (14-25 plants per replicate). Genotypes marked with the same letter did not differ ($P < 0.05$) in Tukey's test.

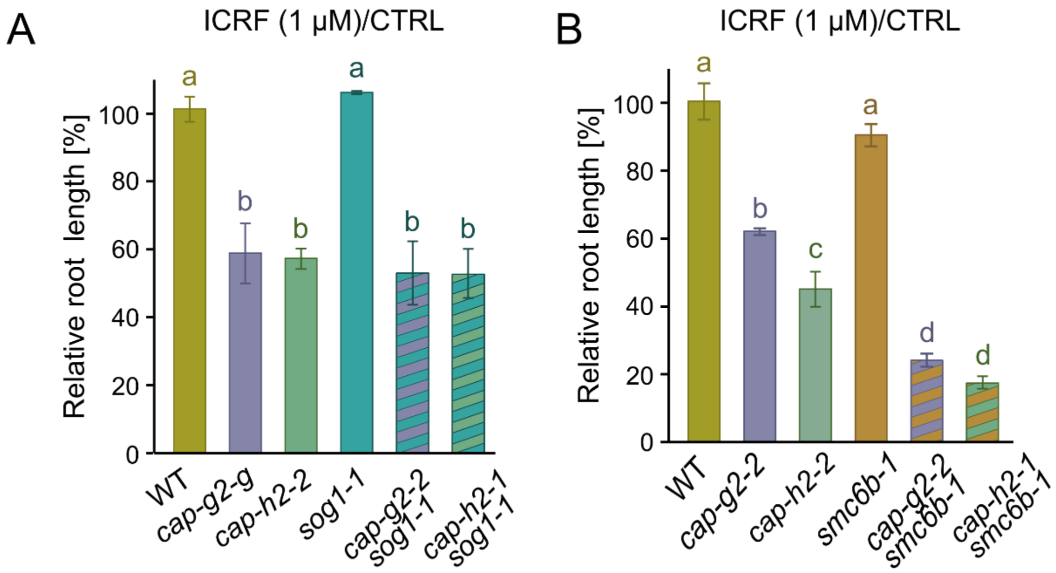


**Figure S9.** Condensin II function is independent of SOG1 and SMC5/6 pathways.

(**A**) Relative root length of seven days old ICRF-187 (ICRF)-treated Condensin II and SOG1 double mutant plants. Error bars are the standard deviation of the means of three biological replicates (14-25 plants per replicate). Genotypes marked with the same letter did not differ ($P < 0.05$) in Tukey's test. The images of representative plants are provided in Figure 5a.

(**B**) Relative root length of Condensin II and SMC6B double mutant plants. The setup and evaluation were done as described in (A). The images of representative plants are provided in Figure 5d.

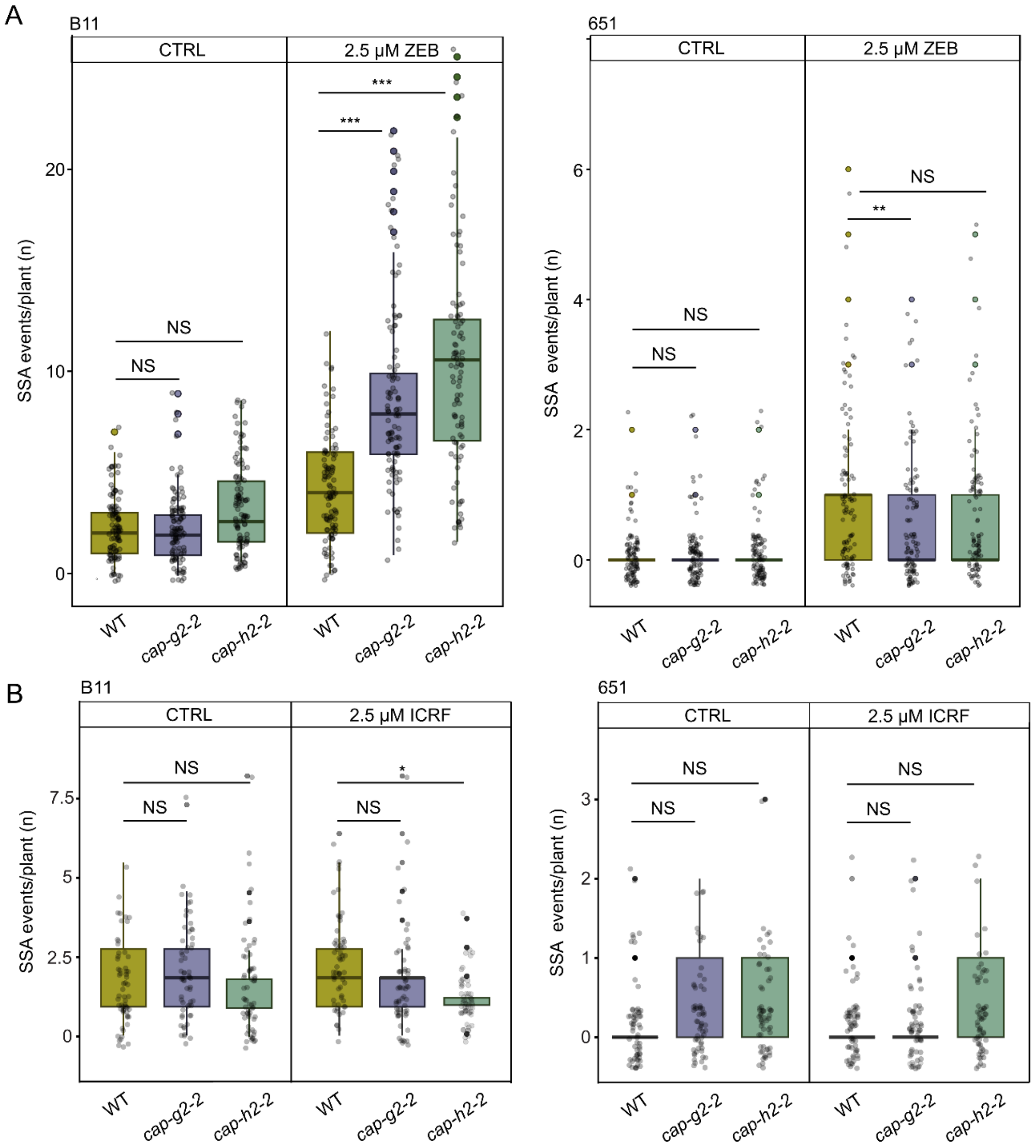


**Figure S10.** Homology-based repair (HBR) in Condensin II mutants.

(**A**) Frequency of single-strand annealing (SSA) events at the reporter loci B11 and 651 in wild type (WT) and Condensin II mutants under control (CTRL) and 2.5 µM zebularine (ZEB) conditions. Individual dots show the number of SSA events per plant. In total 60 – 105 plants per group were evaluated. The boxplots' hinges are in the 1st and 3rd quartile, with a marked median. Whisker marks show the lowest or highest value within the 1.5 interquartile range below or above hinges. Statistical significance was tested with the Kruskall-Wallis H-test with *post hoc* Conover-Iman test of multiple comparisons using rank sums with Benjamini-Hochberg procedure ($P < ½\ \alpha$, $\alpha = 0.05$) in R 4.2.1. *** $P < 0.001$, NS – not significant.

(**B**) Frequency of SSA at the reporter loci B11 and 651 after 2.5 µM ICRF-187 (ICRF) treatment. The parameters were as in (A). In total 60 - 105 plants per time point were evaluated.

**Table S1.** Mutation accumulation experiment. Single nucleotide polymorphisms were identified in the three siblings (R1, R2, R3) of wild-type (WT) and *cap-d3-2* mutant (MUT).

| Geno-type | Position | Ref/ MUT allele | Mutation type | Locus | R1 geno-type | R1 allele depth (ref, alt) | R1 genotype quality | R2 geno-type | R2 allele depth (ref, alt) | R2 genotype quality | R3 geno-type | R3 allele depth (ref, alt) | R3 genotype quality |
|---|---|---|---|---|---|---|---|---|---|---|---|---|---|
| MUT | chr2_4258891 | G/A | Transition | AT2G10820 | A/A | 0,23 | 69 | A/A | 0,22 | 66 | A/A | 0,15 | 45 |
| MUT | chr5_16456485 | C/T | Transition | AT5G41120 | T/T | 0,26 | 78 | T/T | 0,12 | 36 | T/T | 0,26 | 78 |
| MUT | chr5_11141060 | T/C | Transition | AT5G29075 | NA | NA | NA | T/C | 15,7 | 99 | T/C | 15,8 | 99 |
| MUT | chr5_11141073 | A/C | Transversion | AT5G29075 | NA | NA | NA | A/C | 18,8 | 99 | A/C | 16,8 | 99 |
| MUT | chr5_11141078 | A/C | Transversion | AT5G29075 | NA | NA | NA | A/C | 18,8 | 99 | A/C | 15,8 | 99 |
| WT | chr3_14204493 | C/A | Transversion | Intergenic | C/A | 14,30 | 99 | C/A | 19,16 | 99 | C/A | 9,21 | 99 |
| WT | chr3_10753198 | C/T | Transition | Intergenic | C/T | 12,6 | 63 | C/T | 11,11 | 99 | C/T | 14,6 | 38 |
| WT | chr3_2060846 | G/A | Transition | AT3G06610 | G/A | 13,7 | 48 | NA | NA | NA | G/A | 15,7 | 41 |

**Table S2.** Primers used in this study.

| Target | Primer name | Sequence 5' to 3' | Application |
|---|---|---|---|
| *SMC6B* | ET0035 | AGCTTCAACGTGAAATCATGG | genotyping *smc6b-1* |
| *SMC6B* | ET0036 | CTAGACAACATGTCATACCGGG | genotyping *smc6b-1* |
| *T-DNA SALK* | LB_AP1 | ACTGGAACAACACTCAACCCTATCT | genotyping SALK T-DNA lines |
| *SMC2A* | smc2_7F | TTTTGGCTCCACTTTTGTTTG | genotyping *smc2a-1* |
| *SMC2A* | smc2_4R | GATTTGGGATTTCGCTTCTTC | genotyping *smc2a-1* |
| *CAPD2* | LP12 | AAAACCAAGACCATGGAATCC | genotyping *cap-d2-1* |
| *CAPD2* | RP12 | ACACGTGGAGGAAAGTAGGTG | genotyping *cap-d2-1* |
| *CAPH2* | heb2-1_seqF | AGAGATGCTTGTGAATGGAG | genotyping *cap-h2-1* |
| *CAPH2* | heb2-1_seqR | TCAACGTCTCCATTGTTCAC | genotyping *cap-h2-1* |
| *CAPD3* | KP118_cap-d3-2F | TGGTTTGAAAATGGTTGCTTC | genotyping *cap-d3-2* |
| *CAPD3* | KP119_cap-d3-2R | AGCGATAGAAGGAATCGAAGG | genotyping *cap-d3-2* |
| *CAPG2* | KP112_cap-g2-2F | GGTTGCAAATTCAAATGTTCG | genotyping *cap-g2-2* |
| *CAPG2* | KP113_cap-g2-2R | TTCCAATGAGGTCACAAAAGG | genotyping *cap-g2-2* |
| *CAPH2* | KP114_cap-h2-2F | TTTCCGCTCTCTTCAACAGTC | genotyping *cap-h2-2,* cDNA analyses |
| *CAPH2* | KP115_cap-h2-2R | AAAAAGATTGGATGGAGCATTAC | genotyping *cap-h2-2,* cDNA analyses |
| *recB* | B11Fw | TGCTGGTGAACACGTAAAGC | genotyping recB (B11) |
| *recB* | B11Rev | CAGTCGGATGGTTCGTTTCT | genotyping recB (B11) |
| *recA* | recA_F | GAGAAACTTAGCTGGTTGTGATGAT | genotyping recA (651) |
| *recA* | recA_R | CTTTAACCATTGGTCACACTCTCTT | genotyping recA (651) |
| *GUS* | GUSFw | TGGATCGCGAAAACTGTGGA | genotyping recB, recA |
| *GUS* | GUSRev | CGGTGATATCGTCCACCCAG | genotyping recB, recA |
| *SMC2A* | 9A-2_Cf1 | TCGTAAGGATGGAAACGGATC | cDNA analyses *hze4-1* |
| *SMC2A* | 9A-2_Cf2 | CGAGTGAGGGAAATGAGCT | cDNA analyses *hze4-1* |
| *CAPD3* | zdc1 | CTAGCTTACAACAGTTTCGTG | cDNA analyses *hze5-4* |
| *CAPD3* | zdc2 | TCTTGTAGTCTTTGAGGAGGAC | cDNA analyses *hze5-4* |

**Table S3.** P values of the Chi-squared tests were used to assess the statistical relevance of data presented in Figure 6B.

| Chi-squared test P-values for Figure 6B | | | | | | | | |
|---|---|---|---|---|---|---|---|---|
| **Sample** | **WT MOCK** | | | | | | | |
| **WT ZEB** | 0,206 | **WT ZEB** | | | | | | |
| **WT ICRF** | 0,001 | 0,022 | **WT ICRF** | | | | | |
| ***cap-d3-2* MOCK** | 0,000 | | | ***cap-d3-2* MOCK** | | | | |
| ***cap-d3-2* ZEB** | | 0,000 | | 0,111 | ***cap-d3-2* ZEB** | | | |
| ***cap-d3-2* ICRF** | | | 0,000 | 0,004 | 0,077 | ***cap-d3-2* ICRF** | | |
| ***cap-g2-2* MOCK** | 0,000 | | | 0,370 | | | ***cap-g2-2* MOCK** | |
| ***cap-g2-2* ZEB** | | 0,000 | | | 0,859 | | 0,014 | ***cap-g2-2* ZEB** |
| ***cap-g2-2* ICRF** | | | 0,000 | | | x | 0,000 | 0,067 |

**Table S4.** P values of the Chi-squared tests were used to assess the statistical relevance of data presented in Figure 6C.

| Kruskall-Wallis H-test, *post hoc* Conover-Iman with Benjamini-Hochberg for Figure 6C | | | | | | | | |
|---|---|---|---|---|---|---|---|---|
| **Sample** | **WT MOCK** | | | | | | | |
| **WT ZEB** | 0.2490 | **WT ZEB** | | | | | | |
| **WT ICRF** | 0.0585 | 0.1988 | **WT ICRF** | | | | | |
| ***cap-d3-2* MOCK** | 0.0000* | 0.0000* | 0.0000* | ***cap-d3-2* MOCK** | | | | |
| ***cap-d3-2* ZEB** | 0.0000* | 0.0000* | 0.0000* | 0.0224* | ***cap-d3-2* ZEB** | | | |
| ***cap-d3-2* ICRF** | 0.0000* | 0.0000* | 0.0000* | 0.4820 | 0.0320 | ***cap-d3-2* ICRF** | | |
| ***cap-g2-2* MOCK** | 0.0000* | 0.0000* | 0.0000* | 0.3677 | 0.0086* | 0.4085 | ***cap-g2-2* MOCK** | |
| ***cap-g2-2* ZEB** | 0.0000* | 0.0000* | 0.0000* | 0.0000* | 0.0201* | 0.0001* | 0.0000* | ***cap-g2-2* ZEB** |
| ***cap-g2-2* ICRF** | 0.0000* | 0.0000* | 0.0000* | 0.0225* | 0.4783 | 0.0324 | 0.0085* | 0.0165* |

**Table S5.** P values of the Chi-squared tests were used to assess the statistical relevance of data presented in Figure 6D.

| Kruskall-Wallis H-test, *post hoc* Conover-Iman with Benjamini-Hochberg for Figure 6D | | | | | | | | |
|---|---|---|---|---|---|---|---|---|
| **Sample** | **WT MOCK** | | | | | | | |
| **WT ZEB** | 0.3080 | **WT ZEB** | | | | | | |
| **WT ICRF** | 0.0020* | 0.0068* | **WT ICRF** | | | | | |
| ***cap-d3-2* MOCK** | 0.0000* | 0.0000* | 0.0000* | ***cap-d3-2* MOCK** | | | | |
| ***cap-d3-2* ZEB** | 0.0000* | 0.0000* | 0.0000* | 0.0855 | ***cap-d3-2* ZEB** | | | |
| ***cap-d3-2* ICRF** | 0.0000* | 0.0000* | 0.0000* | 0.0000* | 0.0007* | ***cap-d3-2* ICRF** | | |
| ***cap-g2-2* MOCK** | 0.0000* | 0.0000* | 0.0001* | 0.2880 | 0.0277 | 0.0000* | ***cap-g2-2* MOCK** | |
| ***cap-g2-2* ZEB** | 0.0000* | 0.0000* | 0.0000* | 0.0187* | 0.3055 | 0.0015* | 0.0041* | ***cap-g2-2* ZEB** |
| ***cap-g2-2* ICRF** | 0.0000* | 0.0000* | 0.0000* | 0.0000* | 0.0005* | 0.4107 | 0.0000* | 0.0011* |
